# Neural function of Netrin-1 in precancerous lesions of the pancreas

**DOI:** 10.1101/2025.05.14.654046

**Authors:** Hiba Haidar, Anaïs Bellon, Karen Sleiman, Mélanie Hocine, Nicolas Rama, Nicolas Gadot, Darren Carpizo, Patrick Mehlen, Fanny Mann

## Abstract

The nervous system undergoes dynamic structural remodeling to infiltrate cancerous tumors, contributing to their growth and progression. Emerging evidence indicates that neuroplasticity initiates early, with nerve terminals detecting and responding to tissue changes even during precancerous stages. Notably, dense sympathetic axon sprouting has been observed around pancreatic intraepithelial neoplasia (PanIN), a common precursor lesion to pancreatic cancer. However, the molecular signals driving this early neuroplasticity and its functional consequences remain poorly understood. Here, we identify the axon guidance molecule Netrin-1 as a key factor secreted by pancreatic cells within precursor lesions of pancreatic cancer. Netrin-1 promotes sympathetic axon growth and branching through its receptor, Deleted in Colorectal Cancer (DCC). Inhibition of Netrin-1 disrupts sympathetic axon remodeling while accelerating PanIN formation and progression, driven by increased precancerous cell proliferation. Furthermore, human pancreatic tissue analysis corroborates Netrin-1 expression in precursor lesions. These findings suggest that Netrin-1-driven sympathetic neuroplasticity plays a protective role in the precancerous microenvironment by modulating local cellular dynamics, providing new insights into early cancer progression.

## Introduction

The interaction between the nervous system and cancer is bidirectional. The nervous system can influence the initiation and progression of cancer in various tissues, both within and outside the brain. Conversely, cancer can induce profound changes in the structure, function, and connectivity of neural networks. For instance, tumors located in peripheral organs can stimulate nearby nerve terminals to extend and branch deep into their microenvironment^1^. Emerging evidence indicates that this neuroplastic remodeling is not unique to cancer and may begin before malignancy is fully established. In fact, cancers typically do not arise *de novo* but rather progress gradually from premalignant lesions. Studies in organs such as the prostate and pancreas have shown that premalignant lesions already have abnormal patterns of innervation compared to healthy tissues^2–6^.

Pancreatic intraepithelial neoplasia (PanIN) is the most common precursor lesion of pancreatic ductal adenocarcinoma (PDAC). In a transgenic mouse model that spontaneously develops PDAC, we have previously observed that sympathetic axons located in the exocrine pancreas branch out locally and form hotspots of innervation around PanIN lesions ^6^. These findings suggest that sympathetic axon networks reorganize adaptively in response to early tissue changes, with this plasticity potentially playing a regulatory role during the precancerous phase of the disease. To date, the functional role of the sympathetic nervous system in PDAC has been investigated using chemical or surgical nerve ablation. When performed before the onset of malignancy, sympathetic denervation enhances the development and growth of PDAC. This acceleration is caused by a significant increase in protumorigenic CD163-positive macrophages within the tumor microenvironment^6^. These and other findings suggest that the sympathetic nervous system plays a critical role in limiting PDAC progression by modulating immune cell function or recruitment to tissues^6,7^.

The experimental denervation approach eliminates all sympathetic input to the pancreas, offering limited insight into the specific role of disease-related local remodeling of axonal networks surrounding precancerous lesions. A more targeted strategy—such as selectively inhibiting sympathetic axon plasticity while preserving existing nerves—could overcome this limitation. However, the mechanisms driving early pancreatic sympathetic remodeling are still poorly understood. An early study reported a significant increase in the expression of several neurotrophic factors in the pancreas as precancerous lesions develop^4^, but the functional implications of these neurotrophic factors in PanIN innervation remain to be investigated. During the development of axonal projections, families of axon guidance molecules - such as netrins, semaphorins, slits, and ephrins - are critical for the precise navigation of sympathetic axons to their synaptic targets^8^. Although their expression is greatly reduced in adult tissues, expression of axon guidance molecules is reactivated in several pathological conditions. In particular, axon guidance pathway genes are among the earliest and most common mutation targets in pancreatic cancer^9,10^. Extensive research has elucidated some of the functions and mechanisms by which axon guidance molecules influence the tumor ecosystem, including the modulation of cancer cell behavior, such as cancer survival or phenotypic plasticity, cancer-associated fibroblasts, immune cell infiltrates, and angiogenesis^11–13^. However, the potential role of axon guidance molecules in regulating axonal plasticity in the precancerous state remains to be determined.

In this study, we show that the axon guidance molecule Netrin-1 is expressed in precursor lesions of pancreatic cancer and promotes sympathetic axon growth and branching via the Deleted in Colorectal Cancer (DCC) receptor. We also show that inhibition of Netrin-1 function prevents local remodeling of sympathetic axons and accelerates the development of PanINs. This effect is associated with increased proliferation of PanIN cells, but occurs independently of changes in macrophages within the lesion microenvironment. These findings reveal that hyperinnervation of precancerous pancreatic lesions relies on the reactivation of a developmental program for axon growth and guidance and suggest a previously unrecognized protective role for neuroplasticity in the development of precancerous lesions.

## Results

### Neuronal DCC is Essential for Axon Remodeling in Metaplastic Pancreatic Lesions

Sympathetic innervation of the pancreas originates from neurons in the celiac and superior mesenteric ganglia (CSMG), which are located in close proximity to the pancreas. In a healthy mouse pancreas, incoming sympathetic nerve bundles divide into multiple branches that innervate the entire organ, including the exocrine tissue, where their terminals form a network of branching and anastomosing processes (Fig. S1a-e). To investigate how sympathetic terminal innervation is affected by pancreatic acinar injury, we first used a model of chronic pancreatitis. In this model, repeated cerulein injections over 3.5 weeks induced persistent pancreatic inflammation leading to acinar-to-ductal metaplasia (ADM) (Fig. 1a, b). ADM is a reparative program in which acinar cells transdifferentiate into a duct-like phenotype. However, persistent metaplasia can promote progression toward cancer, making ADM one of the earliest precursor lesions in the development of PDAC. Histologically, inflammation-induced metaplastic pancreatic lesions are characterized by ectopic expression of the ductal gene *SOX9* (Sex-determining region Y [SRY]-box transcription factor 9) and macrophage infiltration^14,15^ (Fig. 1c,d’). These metaplastic lesions also showed an increased density of tyrosine hydroxylase (TH)-positive sympathetic axons, whereas the adjacent histologically asymptomatic tissue showed typical innervation (Fig. 1c–e). There was no difference in sympathetic innervation of metaplastic lesions between male and female mice (Fig. S1f).

**Fig. 1:**
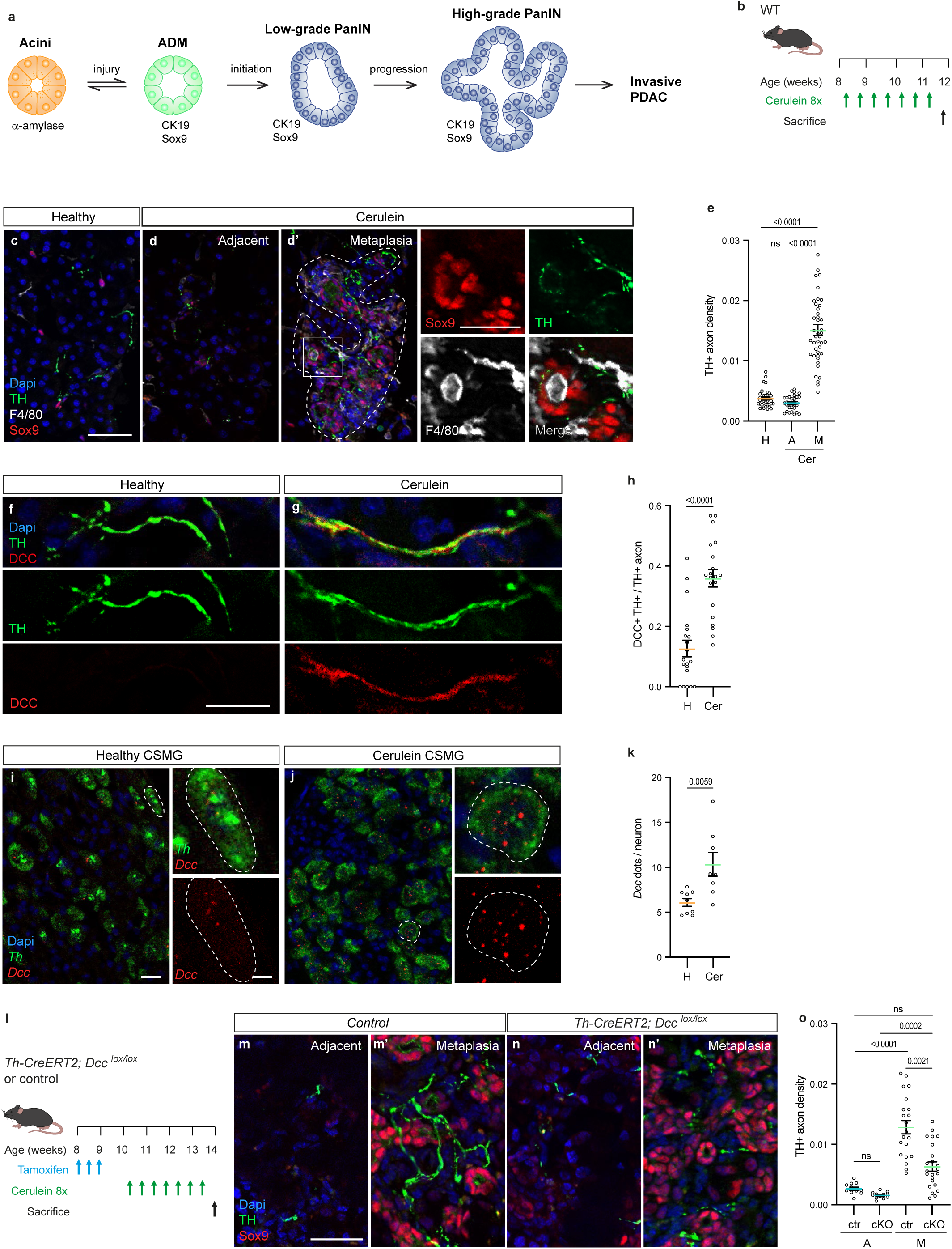
Sympathetic remodeling in metaplastic pancreatic lesions requires DCC. **a**, Illustration showing the progression of pancreatic ductal adenocarcinoma (PDAC). Acinar cells are the main components of the exocrine pancreas responsible for the production of digestive enzymes. Injury or inflammation causes acinar cells to transdifferentiate into duct- like cells by a process called acinar-to-ductal metaplasia (ADM), which can be identified by markers such as *SOX9* or *CK19*. Metaplasia is a reversible process that allows the regeneration of the pancreatic acinar tissue. However, in the presence of oncogenic *Kras* mutations, particularly G12D, ADM progresses to pancreatic intraepithelial neoplasia (PanIN) pre-cancerous lesions. PanINs show varying degrees of neoplasia, ranging from low to high- grade carcinoma in situ. **b**, Protocol for the induction of ADM in chronic pancreatitis induced by repeated cerulein injections. Green arrows indicate the days of cerulein administration, with mice receiving hourly injections eight times on the treatment day. Samples were collected 2 days after the last cerulein dose. **c-d,** Representative immunohistochemistry (IHC) staining for TH (a marker for sympathetic neurons) and F4/80 (a pan-macrophage marker) on sections of the exocrine pancreas from healthy (untreated) mice and cerulein-treated mice. Cerulein-induced metaplasia, identified by *SOX9* staining (d’), showed increased sympathetic innervation and macrophage infiltration compared to the adjacent asymptomatic tissue (d) or healthy pancreas. Insets in d’ provide a zoomed-in view of individual staining. DAPI for nuclear staining. Scale bar: 50 µm, 25 µm (insets). **e**, Quantification of TH^+^ sympathetic axon density in the exocrine pancreas of healthy mice (H), metaplastic lesions (M), and adjacent exocrine tissue (A) of cerulein-treated mice (Cer). Data are expressed as mean ± SEM. *n* = 36 ROIs from 6 healthy mice, 44 metaplastic, and 31 adjacent ROIs from 6 cerulein-treated mice. p values indicated (ns: p > 0.05) (Kruskal–Wallis test). **f-g,** Representative IHC staining for TH and DCC on sections of the exocrine pancreas from healthy and cerulein-treated mice. DAPI for nuclear staining. Scale bar: 20 µm. **h**, Quantification of the ratio of TH^+^ sympathetic axons co-labeled for DCC in healthy (H) and cerulein-treated (Cer) pancreas. Data are presented as mean±SEM. *n*­=20 sections from 3 mice per group. p values indicated (unpaired t-test). **i-j**, Representative RNAscope *in situ* hybridization for *Th* mRNA and *Dcc* mRNA in sections of the celiac superior mesenteric ganglia (CSMG) of healthy (H) and cerulein-treated mice (Cer). Higher magnification insets highlight individual sympathetic neuronal cell bodies. Scale bars: 20 µm and 5 µm (insets). **k**, Quantification of *Dcc* mRNA (dots) in CSMG sympathetic neurons from healthy (H) and cerulein-treated (Cer) mice. Data are presented as mean±SEM. *n*­=9 sections from 3 healthy mice and 8 sections from 3 cerulein-treated mice. p values indicated (unpaired t-test). **l**, Experimental design for tamoxifen injection time points and cerulein treatment used in *Th- CreERT2–Dcc ^lox/lox^* and control *Cre*-negative *Dcc^lox/lox^*mice. **m-n**, Representative IHC staining for TH and *SOX9* in metaplastic pancreatic lesions (m’ and n’) and adjacent exocrine tissue (m and n) of cerulein-treated control and *Th-CreERT2–Dcc ^lox/lox^*mice. DAPI for nuclear staining. Scale bar: 25 µm. **o**, Quantification of TH^+^ sympathetic axon density in metaplastic pancreatic lesions (M) and adjacent tissues (A) of cerulein-treated control (ctr) and *Th-CreERT2–Dcc ^lox/lox^* (cKO) mice. Data are presented as mean±SEM. *n*­=21 metaplastic and 11 adjacent ROIs from 3 control mice; 23 metaplastic and 12 adjacent ROIs from 3 cKO mice. p values indicated (ns: p > 0.05) (Kruskal–Wallis test).

We used the chronic pancreatitis model to investigate the molecules that drive remodeling of sympathetic axon terminals. First, we performed an immunohistochemical screen to profile the expression of receptors for the canonical axon guidance molecules that may mediate intercellular communication between sympathetic axons and their microenvironment. Among these, the DCC receptor showed a particularly interesting profile: while absent from sympathetic axons in the healthy pancreas, DCC was detected on sympathetic axons following cerulein treatment (Fig. 1f–g; Fig. S2a). To rule out the possibility that the presence of DCC was merely due to an increase in axonal density, we quantified the proportion of sympathetic axons expressing DCC and confirmed a true increase in DCC expression within these axons (Fig. 1h). This finding was further validated by RNAscope *in situ* hybridization of sections of the CSMG containing the sympathetic neuronal cell bodies (Fig. 1i–k; Fig. S2b). Importantly, DCC expression was predominantly observed in TH^+^ sympathetic neurons, with no staining observed in other cells in the pancreatic environment (Fig. S2c, d). DCC was notably absent from other axon types innervating the pancreas, such as VAChT^+^ cholinergic axons and CGRP^+^ sensory axons (Fig. S2e-h). Furthermore, sympathetic axons in both healthy and metaplastic pancreases did not express UNC5B—an axon guidance receptor that serves as a non-obligatory co-receptor for DCC (Fig. S2i, j). However, UNC5B was detected in macrophages (Fig. S2k-m), as previously reported^16^.

To investigate the role of DCC in injury-induced sympathetic remodeling, we generated *Th- CreERT2; Dcc^lox/lox^* mice, enabling inducible deletion of *Dcc* in sympathetic neurons, including those residing in the CSMG (Fig. S3a, b). *Dcc* ablation in adult mice did not alter baseline sympathetic innervation of the exocrine pancreas (Fig. S3c–e). However, in the context of cerulein-induced chronic pancreatitis, *Dcc* loss profoundly attenuated sympathetic hyperinnervation within metaplastic lesions, while sparing adjacent tissue innervation (Fig. 1l–o). Like sympathetic axons, VAChT^+^ cholinergic axons exhibited increased density in metaplastic regions, yet their abundance remained unaffected by *Dcc* deletion (Fig. S3f-h). Additionally, CGRP+ axons failed to innervate metaplastic areas under all conditions (Fig. S3i-k). Thus, the loss of *Dcc*-dependent sympathetic remodeling did not trigger compensatory sprouting or recruitment of alternative axon types. These findings establish *Dcc* as a pivotal regulator of lesion-induced sympathetic plasticity in the pancreas.

### Metaplastic Pancreatic Cells Drive Axonal Remodeling via Netrin-1 Signaling

We next examined the expression of the DCC ligand Netrin-1. Although no expression was detected in the healthy pancreas, high levels of Netrin-1 protein were detected in metaplastic pancreatic cells and in infiltrating macrophages in the cerulein-treated pancreas (Fig. 2a–e; Fig. S4a). This up-regulation was confirmed by RNAscope in situ hybridization (Fig. S2b; S4b-d). We next examined the in vitro response of CSMG neurons to Netrin-1. Cultured adult CSMG neurons displayed DCC expression along their extending neurites (Fig. S4e, f) and demonstrated a marked increase in neurite outgrowth upon exposure to recombinant Netrin-1 (Fig. 2f-j). Netrin-1 has two receptor binding sites: a DCC-only binding site and a generic receptor binding site for DCC or UNC5B^17–19^. We observed that the growth-promoting effect of Netrin-1 was efficiently blocked by an anti-Netrin-1 monoclonal antibody (mAb) targeting the generic receptor-binding site, but not by a control isotype antibody^18^ (Fig. 2h–j). The anti-Netrin-1 mAb may work by inhibiting the Netrin-1-induced formation of DCC homodimers, which is the active signaling state of the receptor^17^. We then administered the anti-Netrin-1 mAb to mice *in vivo* during the induction of chronic pancreatitis (Fig. 2k). Analysis of tissue sections and whole-mount cleared pancreas showed that Netrin-1 inhibition significantly reduced sympathetic axon growth and branching within metaplastic lesions (Fig. 2l–o; Fig. S5a–c), without affecting the pre-existing sympathetic network in the adjacent tissues (Fig. S5a–e). VAChT^+^ cholinergic innervation of metaplastic lesions was unaffected by treatment (Fig. S5f-h), and no CGRP^+^ sensory fibres were observed in metaplastic lesions under any conditions (Fig. S5i-k). Similarly, netrin blockade did not affect the density of blood vessels (Fig. S6a-c) or the number and type of macrophages (Fig. S6d-l) present in the lesions.

**Fig. 2:**
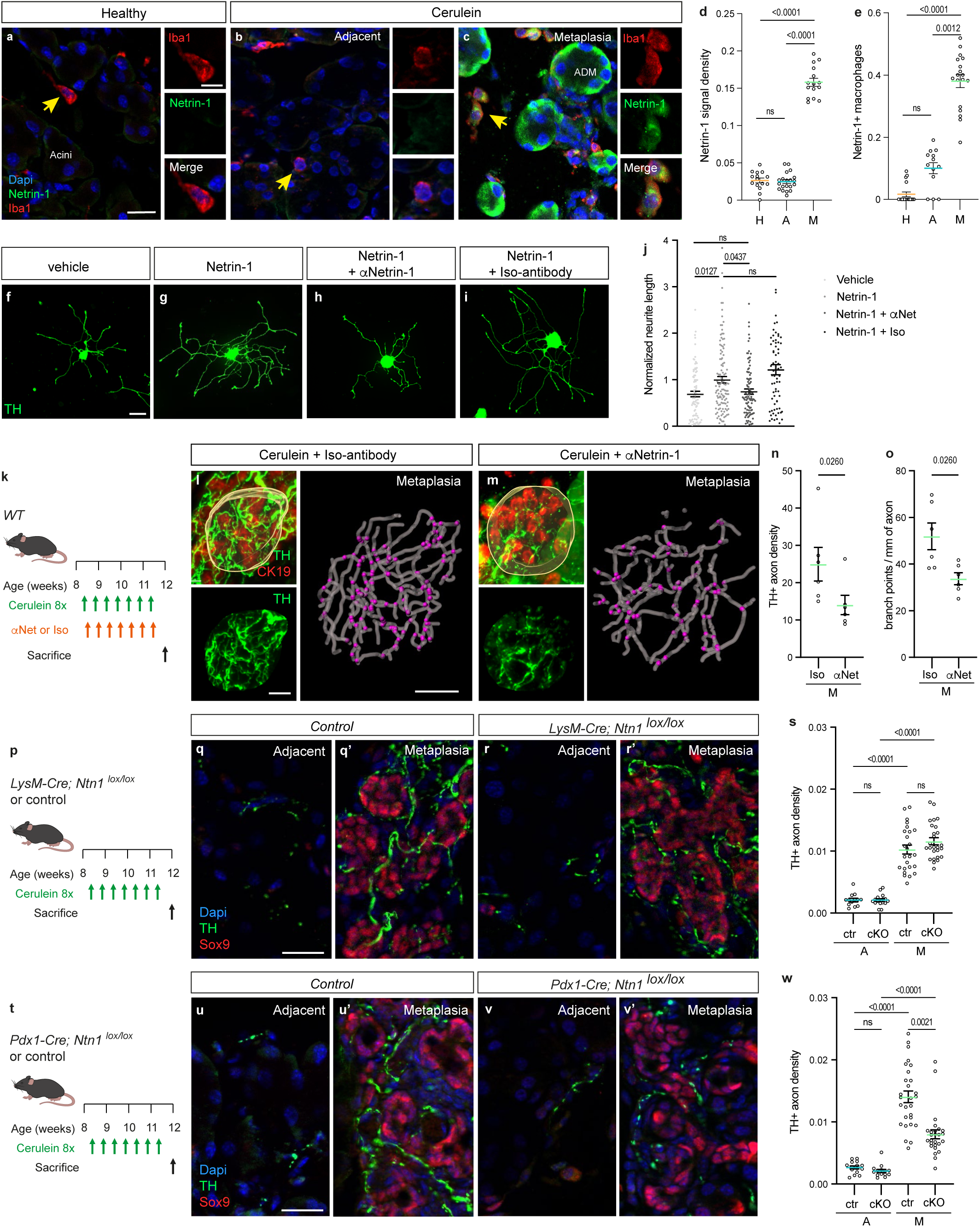
Netrin-1 expression in metaplastic cells induces sympathetic axon remodeling. **a-c**, Representative immunohistochemistry (IHC) staining for Netrin-1 and Iba1 (a pan- macrophage marker) in sections of healthy pancreas (a), and cerulein-treated pancreas (b: adjacent asymptomatic tissue; c: metaplasia). Yellow arrows indicate enlarged macrophages in the insets. DAPI for nuclear staining. Scale bar: 20 µm and 5 µm (inset). **d**, Quantification of Netrin-1 fluorescence labeling in the exocrine pancreas of healthy (H) mice, metaplastic lesions (M) and adjacent exocrine tissue (A) of cerulein-treated mice. Data are presented as mean±SEM. *n* = 15 ROIs from 3 healthy mice, 15 metaplastic, and 20 adjacent ROIs from 3 cerulein-treated mice. p-values are indicated on the graph (Kruskal- Wallis test). **e**, Quantification of the proportion of Iba1^+^ macrophages expressing Netrin-1 in the healthy (H) pancreas, metaplastic lesions (M) and adjacent exocrine tissue (A) of cerulein-treated mice. Data are presented as mean±SEM. *n* = 18 ROIs from 3 healthy mice, 18 metaplastic, and 13 adjacent ROIs from 3 cerulein-treated mice. p-values are indicated on the graph (Kruskal-Wallis test). **f-i**, Representative images of TH^+^ sympathetic neurons from the CSMG cultured in PBS (vehicle) or recombinant Netrin-1 protein, together with an anti-Netrin-1 blocking antibody (αNetrin-1) or an isotype control antibody (Iso-antibody). Scale bar: 50 μm. **j**, Quantification of total neurite length in cultures shown in (f–i). Data are presented as mean±SEM and normalized to 1 for the Netrin-1 condition. *n*­=94 (vehicle), 132 (Netrin-1), 94 (Netrin-1 + αNetrin-1), and 89 (Netrin-1 + Iso-antibody) neurons from four independent experiments. p values indicated (ns: p > 0.05) (Kruskal–Wallis test). **k**, Experimental design for cerulein injections and subsequent treatments with αNetrin-1or Iso-antibody. **l-m**, Maximum intensity projection of 3D images of TH^+^ sympathetic axon networks in CK- 19+ metaplastic pancreatic lesions treated with Iso-antibody (l) or αNetrin-1 (m). Right panels show 3D reconstruction of sympathetic axons and branching points (purple dots) within the yellow-segmented volume. Scale bar: 30 µm. **n-o**, 3D quantification of TH^+^ sympathetic axon density (n) and branching (o) in metaplastic lesions (M) treated with αNetrin-1 (αNet) or Iso-antibody (Iso). Data are presented as mean±SEM. *n*­=6 ROIs from 3 mice per condition. p-values indicated (Mann–Whitney test). **p**, Experimental design for cerulein treatment of *LysM-Cre–Ntn1^lox/lox^* and control mice. **q-r**, Representative IHC staining for TH and *SOX9* in metaplastic lesions (q’ and r’) and adjacent tissue (q and r) of cerulein-treated *LysM-Cre–Ntn1^lox/lox^* and control mice. DAPI for nuclear staining. Scale bar: 25 µm. **s**, Quantification of TH^+^ sympathetic axon density in metaplastic pancreatic lesions (M) and adjacent tissues (A) of cerulein-treated control (ctr) and *LysM-Cre–Ntn1^lox/lox^* (cKO) mice. Data are presented as mean±SEM. *n*­=25 metaplastic and 13 adjacent ROIs from 3 control mice; 26 metaplastic and 16 adjacent ROIs from 3 cKO mice. p values indicated (ns: p > 0.05) (Kruskal–Wallis test). **t**, Experimental design for cerulein treatment of *Pdx1-Cre–Ntn1^lox/lox^* and Cre-negative control mice. **u-v**, Representative IHC staining for TH and Sox9 in metaplastic pancreatic lesions (u’ and v’) and adjacent exocrine tissue (u and v) of cerulein-treated control and *Pdx1-Cre–Ntn1^lox/lox^*mice. DAPI for nuclear staining. Scale bar: 25 µm. **w**, Quantification of TH^+^ sympathetic axon density in metaplastic pancreatic lesions (M) and adjacent tissues (A) of cerulein-treated control (ctr) and *Pdx1-Cre–Ntn1^lox/lox^* (cKO) mice. Data are presented as mean±SEM. *n*­=28 metaplastic and 15 adjacent ROIs from 3 control mice; 26 metaplastic and 14 adjacent ROIs from 3 cKO mice. Significant p values are indicated on the graph (ns: p > 0.05) (Kruskal–Wallis test).

To identify the relevant cellular source of Netrin-1 for sympathetic axon remodeling, we selectively silenced the loxP-flanked *Ntn1* allele in either macrophages (using LysM-Cre-mediated recombination)^20,21^ or in pancreatic cells (using Pdx1-Cre-mediated recombination) (Fig. S7a–c). In unchallenged conditions, neither *LysM-Cre–Ntn1^lox/lox^* nor *Pdx1-Cre– Ntn1^lox/lox^* mice exhibited significant alterations in pancreatic structure or sympathetic innervation (Fig. S7d–i). However, after cerulein treatment, the loss of Netrin-1 in pancreatic cells, but not in macrophages, markedly suppressed sympathetic axon sprouting within metaplastic lesions (Fig. 2p–w). These findings reveal that Netrin-1, secreted by metaplastic acinar cells reprogrammed into duct-like cells, drives sympathetic neuroplasticity by activating its DCC receptor on neighboring neurons.

### Blocking Netrin-1 disrupts PanIN innervation and progression

Next, to test the importance of the Netrin-1/DCC signaling pathway in precancer-associated neuroplasticity, we used transgenic KIC (*Kras^LSL-G12D/+^; InkA4/Arf^lox/lox^; Pdx1-Cre*) mice. These mice spontaneously develop pancreatic cancer, with a progression from low-grade to high-grade PanINs, ultimately advancing to PDAC (Fig. 1a). We observed high levels of Netrin-1 expression in PanIN precursor lesions compared to adjacent asymptomatic tissues at both protein (Fig. 3a-d) and RNA levels (Fig. S8a-c). In contrast, there was little or no detectable expression of Netrin-1 in the pancreatic cancer cells of the KIC PDAC (Fig. 3c-d), which is consistent with a previous report (Dudgeon et al., 2023). DCC was upregulated in sympathetic neurons of the CSMG of KIC mice (Fig. 3e-g). As the KIC transgenic model limits other genetic manipulations, we used the anti-Netrin-1 mAb to block Netrin/DCC signaling. KIC mice were treated with the antibody from 4 to 6.5 weeks of age, corresponding to the onset and progression of the precancerous phase in this model (Fig. 3h). After treatment, whole-mount pancreases were immunostained for TH to label sympathetic neurons, and PanINs were identified either by tissue autofluorescence or by co-staining with anti-cytokeratin 19 antibody ^6^. As previously reported, PanIN lesions were associated with areas of sympathetic hyperinnervation characterized by increased axon density and branching^6^ (Fig. 3i-n). Following Netrin-1 blockade, a significant reduction in sympathetic innervation of PanINs was observed, while the normal pattern of innervation in adjacent tissues remained unchanged (Fig. 3k-n). Thus, Netrin-1 is a key driver of precancerous innervation.

**Fig. 3:**
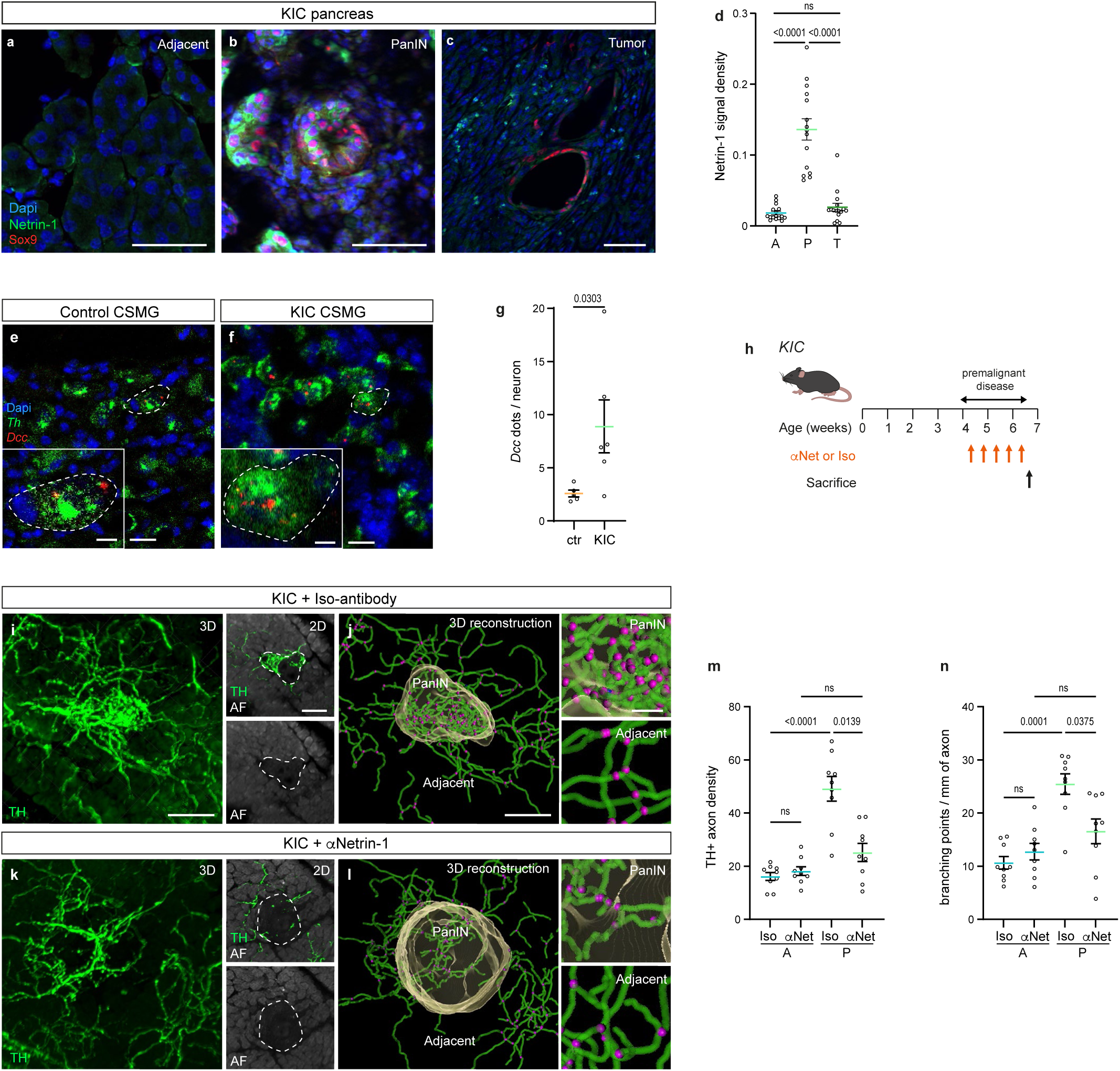
Netrin-1 promotes the innervation of precancerous PanIN lesions. **a-c**, Representative immunohistochemistry staining for Netrin-1 and Sox9 in pancreatic sections through PanIN (b), PDAC tumor (c) and adjacent asymptomatic tissue (a) obtained from 6.5-week-old KIC mice. DAPI nuclear staining. Scale bar: 50 µm. **d**, Quantification of Netrin-1 fluorescence labeling in PanIN (P), PDAC tumor (T), and adjacent tissue (A). Data are presented as mean±SEM. *n*­=15 PanINs, 16 PDAC, and 16 asymptomatic ROIs from 3 KIC mice. p-values are indicated on the graph (Kruskal–Wallis test). **e-f**, Representative RNAscope *in situ* hybridization for *Th* mRNA and *Dcc* mRNA in sections of the celiac superior mesenteric ganglia (CSMG) from 6.5-week-old control (e) and KIC (f) mice. Higher magnification insets highlight individual sympathetic neuronal cell bodies. Scale bars: 20 µm and 5 µm (insets). **g**, Quantification of *Dcc* mRNA (dots) in CSMG sympathetic neurons from 6.5-week-old control (ctr) and KIC mice. Data are presented as mean±SEM. *n*­=5 CSMG sections from 3 control mice and 6 CSMG sections from 3 KIC mice. P-values are indicated on the graph (Mann–Whitney test). **h**, Experimental design for the treatment of KIC mice with either anti-Netrin-1 antibody (αNetrin-1) or isotype control antibody (Iso-antibody). **i-l**, left panels in (i) and (k) show maximum intensity projection of 3D images of TH^+^ sympathetic networks in PanIN and adjacent tissue treated with Iso-antibody (i) or αNetrin-1 (k) antibodies. Right panels in (i) and (k) show 2D optical sections of PanINs identified by tissue autofluorescence (AF). (j) and (l) show the 3D reconstruction of sympathetic axons and branching points (purple dots) in PanINs and adjacent tissues. The volume of the PanIN is shown in yellow. Scale bars: 100 µm (i, k, and left panels of j and l), 20 µm (right panels of j and l). **m-n**, 3D quantification of TH^+^ sympathetic axon density (m) and branching (n) in PanIN (P) and adjacent (A) tissues of KIC mice treated with Iso-antibody (Iso) or αNetrin-1 (αNet). Data are presented as mean±SEM. *n*­=6 PanINs and 6 asymptomatic ROIs from 3 mice per group. Significant p-values are indicated on the graphs (ns: p > 0.05) (Kruskal–Wallis test).

In our previous work, we showed that sympathetic denervation of the pancreas significantly upregulated the pro-tumorigenic macrophage marker CD163 in tumors of KIC mice^6^. Building on this, we investigated whether blocking local sympathetic remodeling in PanINs using anti-Netrin-1 antibody would have a similar effect. However, we observed no significant changes in macrophage density or subtypes, including CD163-positive macrophages (Fig. S8d-l). These results suggest that local innervation does not directly regulate macrophage recruitment or polarization in the PanIN microenvironment.

Despite the lack of effect on macrophages, disruption of local axonal remodeling was associated with an increased number of PanIN foci (Fig. 4a-c) and a higher proportion of high-grade compared to low-grade PanIN lesions (Fig. 4d-f). The pro-PanIN effect was attributed to increased proliferation (Ki67-positive; Fig. 4g-i) of duct-like cells without affecting apoptosis (cleaved caspase-3-positive) (Fig. S8m-o). Collectively, these findings suggest a role for sympathetic axonal remodeling as a protective mechanism that attenuates the progression of precancerous lesions (Fig. 4j).

**Figure 4:**
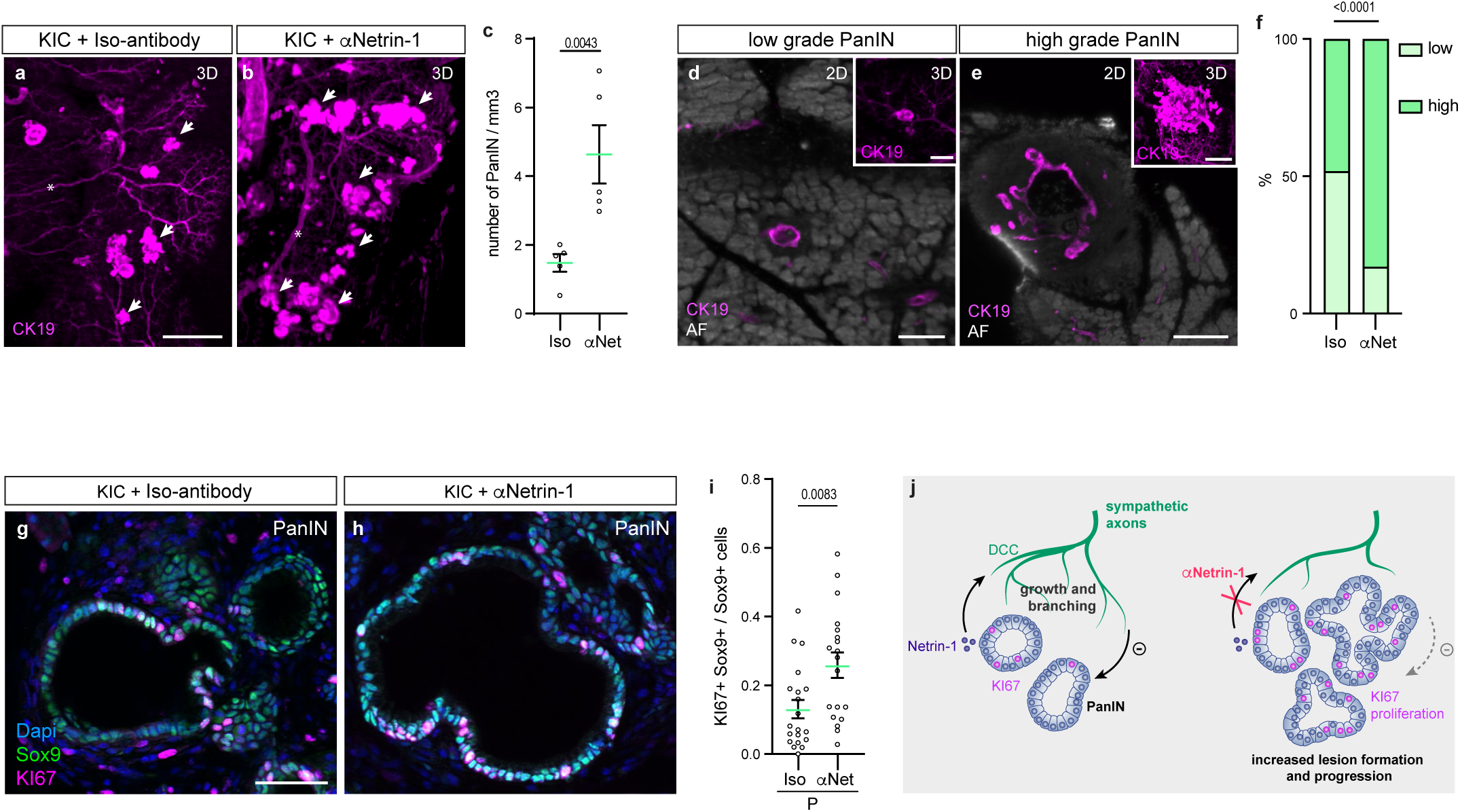
Netrin-1 signaling inhibits the progression of precancer PanIN lesions. **a-b**, Maximum intensity projection of 3D images of CK19 immunostaining in KIC mice treated with anti-Netrin-1 antibody (αNetrin-1) or isotype control antibody (Iso-antibody). CK19 labels both pancreatic ducts and their ramifications (asterisks), and PanIN lesions (arrows). Scale bar: 500 µm. **c**, Quantification of the number of PanIN lesions per mm^3^ in KIC mice treated with Iso- antibody (Iso) or αNetrin-1 (αNet). Data are presented as mean±SEM. *n*­=6 ROIs (Iso) and 5 ROIs (αNet) from 3 mice per group. p-values are indicated on the graph (Mann– Whitney test). **d-e**, 2D optical sections illustrating low-grade (d) and high-grade (e) PanIN lesions. PanINs were labeled with CK19, and pancreatic tissue was visualized by autofluorescence (AF). The insets show the maximum intensity projections of 3D CK19^+^ signals. Scale bars: 100 µm (d), 200 µm (e). **f**, Proportions of low- and high-grade PanIN lesions in KIC mice treated with Iso-antibody (Iso) or αNetrin-1 (αNet). *n*­=20 (Iso) and 41 (αNet) PanINs from 3 mice per group. Data are presented in percentage. p-values are indicated on the graph (Chi-squared test). **g-h**, Representative immunohistochemistry staining for Sox9 and KI67 in pancreatic sections through PanIN lesions in KIC mice treated with Iso-antibody (g) or αNetrin-1 (h). DAPI for nuclear staining. Scale bar: 50 µm. **i**, Quantification of the proportion of Sox9+ cells expressing KI67 in PanIN lesions in KIC mice treated with Iso-antibody (Iso) or αNetrin-1 (αNet). Data are presented as mean ± SEM. *n* = 20 (Iso) and 19 (αNet) ROIs from 3 mice per group. p-value is indicated on the graph (Mann–Whitney test). **j**, Summary of findings and proposed model: Reactivation of the neurodevelopmental Netrin- 1/DCC signaling pathway promotes hyperinnervation of non-malignant pancreatic lesions by sympathetic nerve terminals. This process may initiate a negative feedback loop that restrains lesion progression to higher grades by limiting cell proliferation.

### Netrin-1 is expressed pancreatic lesions in human

The above results show that pancreatic metaplastic and precancerous cells secrete Netrin-1 to recruit local sympathetic nerve endings. To confirm this in a human context, we analyzed Netrin-1 expression in the human pancreas by re-examining single nucleus RNA sequencing (sNuc-seq) data from chronic pancreatitis samples from two human donors ^22^. Consistent with the altered cellular composition observed in chronic inflammation, acinar cells were largely outnumbered by two populations of duct-like cells (Duct-like 1 and Duct-like 2) and chemosensory tuft cells were identified, which also arise from acinar cell plasticity (Fig. 5a and S9a). NTN1 was not expressed in acinar cells but was present in both duct-like populations, with the highest expression levels in the Duct-like 1 cluster, as well as in tuft cells (Fig. 5b-c and S9b). To further extend these findings, we performed immunohistochemistry for Netrin-1 on tissue sections from three different donor pancreata. No significant expression of Netrin-1 was detected in normal pancreatic tissue (Fig. 5d). However, in sections from tissue adjacent to PDAC, Netrin-1 staining was significantly elevated in areas of ADM and PanIN lesions (Fig. 5e-h). These results demonstrate that Netrin-1 is upregulated in precursor lesions of PDAC in humans, similar to observations in mice.

**Fig. 5:**
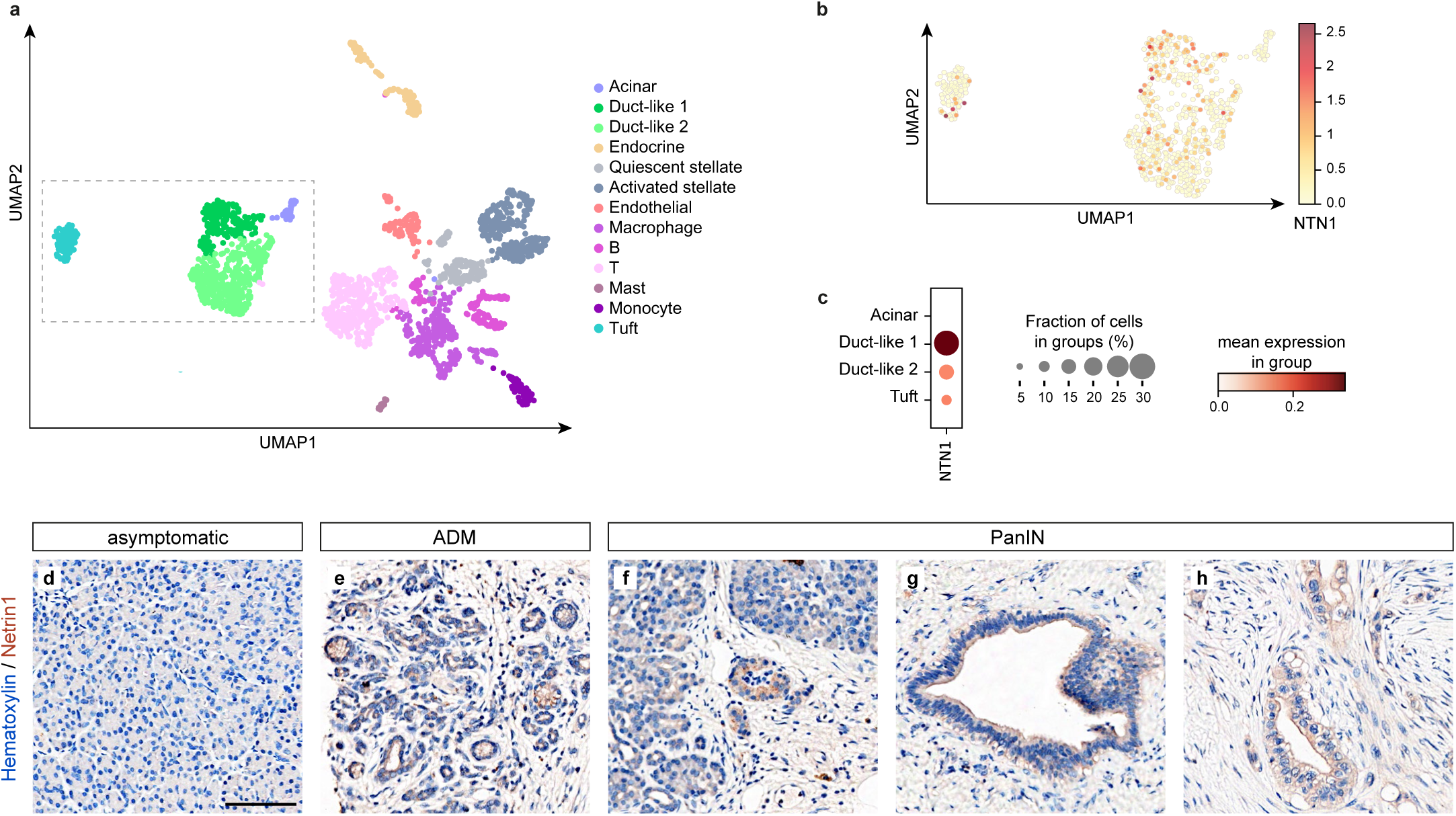
Netrin-1 expression in ADM and PanIN lesions in the human pancreas. **a**, Uniform Manifold Approximation and Projection (UMAP) of all cells from single-nucleus RNA sequencing of two donors with chronic pancreatitis, with populations identified by color. **b**, UMAP plot showing NTN1 expression in acinar, ductal, and tuft cell clusters. **c**, Dot plot of NTN1 expression in acinar, ductal, and tuft cell clusters, where dot size represents the percentage of expressing cells and color indicates average expression levels. **d-h**, Immunohistochemical analysis of Netrin-1 expression in histological sections from human PDAC patients, counterstained with hematoxylin. Normal (asymptomatic) tissue (d) and PanIN (f) were from a stage IB patient, while ADM (e) and PanIN (g,h) from peritumor tissues of stage III patients. Scale bar: 100 µm.

## Discussion

During development, Netrin-1/DCC signaling orchestrates the coordinated patterning of the nervous and vascular systems by guiding sympathetic innervation of blood vessels—a key mechanism for regulating organ physiology via vascular control ^23^. Here, we show that this axon guidance pathway is co-opted in early pancreatic lesions to support sympathetic axon remodeling and provide protection against precancer development.

Netrin-1 is frequently overexpressed in cancerous tissue. This study reveals its involvement in precancer by showing its activation in the pancreas during metaplastic transition, when acinar cells reprogram toward an epithelial-like phenotype. Netrin-1 expression is notably higher in precursor lesions of pancreatic cancer, such as ADM and PanIN, in both murine and human tissues, compared to levels observed in PDAC. The mechanisms driving this early induction remain unknown. During metaplastic transition, cells undergo extensive epigenetic changes, which which may unlock developmental programs normally active in the embryonic pancreatic epithelium (as is the case for Netrin-1,^24^), but silenced in adult tissues.

This study demonstrates that inhibiting Netrin-1 function selectively reduces sympathetic hyperinnervation in precursor lesions of pancreatic cancer, with no effect on innervation in healthy tissue. In a pulmonary fibrosis model, Netrin-1 expression by macrophages has been shown to promote sympathetic axon remodeling^25^. However, in the pancreas, the principal source of Netrin-1 is pancreatic cells themselves, as deletion of Netrin-1 in macrophages had no significant effect on innervation. Given that Netrin-1 can act as an angiogenic cue^26^, it was important to consider whether its effects on axon growth might be mediated indirectly via vascular remodeling, since blood vessels are normally innervated by sympathetic axons. However, inhibition of Netrin-1 did not alter blood vessel density in pancreatic lesions, although the functionality of these vessels has not been assessed. This suggests that blood vessels do not play a substantial role in axon sprouting. This finding aligns with previous studies showing that axonal and vascular networks remodel in parallel but remain largely non-overlapping in pancreatic lesions^6^. Together, these results support a direct role of Netrin-1 secretion by metaplastic and precancerous cells in promoting the growth of neighboring sympathetic axons. Consistent with this, deletion of the Netrin-1 receptor DCC specifically in sympathetic neurons abolished lesion-induced sympathetic axon outgrowth.

The tumor-promoting role of Netrin-1 has been well established in multiple cancer models, where it often confers a selective survival advantage to cancer cells, enhancing their ability to form aggressive tumors. In PDAC, Netrin-1 has been shown to drive tumor progression and facilitate liver metastasis^27,28^. This effect is independent of the nervous system and occurs via an autocrine mechanism, whereby Netrin-1 inhibits the pro-apoptotic activity of the Unc5b receptor in cancer cells. In contrast, this mechanism does not appear to be active in metaplastic or precancerous pancreatic cells, which lack Unc5 receptor expression and do not exhibit increased apoptosis upon Netrin-1 inhibition.

In contrast to its protumorigenic role in cancer, Netrin-1 functions as a protective factor during precancerous stages by limiting PanIN formation and halting progression to higher-grade lesions. This protective effect is likely mediated through enhanced sympathetic innervation. This aligns with previous studies in KIC mice, where global sympathetic denervation performed prior to lesion onset revealed a protective role for the sympathetic nervous system^6^. However, despite achieving similar outcomes, global denervation and local inhibition of sympathetic remodeling appear to act via distinct mechanisms. Specifically, sympathetic denervation regulates the recruitment of CD163-positive macrophages in the diseased pancreas^6^, whereas inhibition of local sympathetic remodeling — which preserves existing sympathetic networks — does not affect CD163-positive or other macrophage subtypes. These observations suggest that the sympathetic nervous system exerts dual and context-dependent roles: at the systemic and organ levels, it regulates immune cell activation and tissue infiltration, thereby preventing excessive immune suppression; at the lesion level, it exerts a negative regulatory influence on the proliferative behavior of precancerous cells. Whether this local effect involves direct neural signaling remains to be determined, as adrenergic receptor activation has been shown to suppress proliferation in certain cell types^29–31^. Finally, other studies have reported a tumor-promoting role for the sympathetic nervous system in PDAC. This discrepancy may reflect differences in timing and experimental models, as those studies were typically conducted when malignant tumors were already present, or used cancer cell transplantation models - often in immunodeficient mice - that bypass the early, precancerous phase^32,33^.

Anti-Netrin-1 therapy is currently being evaluated in clinical trials for the treatment of several cancers, including PDAC, with promising early results ^34^. Our findings suggest that such treatment may also impact precancerous PanIN lesions, which are frequently located at the periphery of the tumor. The potential progression of these lesions under anti-Netrin1 therapy will therefore warrant careful monitoring. Recent studies have also identified PanINs - sometimes numbering in the hundreds - in individuals without overt pancreatic pathology^35^. Based on our work, Netrin-1 could serve as a potential biomarker for precancerous lesions ^36^, offering a tool for surveillance of their evolution in individuals at high risk of developing pancreatic cancer.

## Methods

### Mice

All animal experiments were performed in compliance with the guidelines of the French Ministry of Agriculture (agreement number F1305521) and were approved by the Ethics Committee for Animal Experimentation of Marseille–CEEA-014 under approval numbers APAFIS#27289-2020091713522725 v4 and APAFIS #49983-2024092515422118 v1. Tumor burden was assessed using a body condition score and the maximum tumor burden allowed by the Ethics Committee was not exceeded in all experiments. Mice were group-housed (no more than 5 mice per cage) in individually ventilated cages under controlled conditions (room temperature [RT]: 23 ± 2°C; 12-h light/dark cycle; humidity: 45–65%) in a pathogen-free facility, and were given unrestricted access to food and water. C57BL/6 mice obtained from Janvier Laboratories (Le Genest-Saint-Isle, France) were used as the wild-type (WT) strain. *Ntn1^lox/lox^* mice ^37^ were crossed with either *LysM-Cre* ^38^ (JAX stock #004781) or *Pdx1-Cre* ^39^ mice to delete Netrin-1 in macrophages and pancreatic cells, respectively. *Th-CreERT2* mice^40^ (JAX stock #025614) were crossed with *Dcc^lox/lox^*^41^ to induce the deletion of DCC expression in adult sympathetic neurons, and with the *R26R-EYFP* reporter mice ^42^(JAX stock #006148) to report Cre activity after tamoxifen treatment. *S100b-EGFP* mice ^43^ were used for dissection of the celiac superior mesenteric ganglia (CSMG) for culturing. KIC mice were obtained by crossing *Kras^LSL-G12D^*mice ^44^, *Ink4a/Arf ^lox/lox^*^45^, and *Pdx1-Cre* mice. All transgenic animals were maintained on a C57BL/6 background. Experiments were performed on adult (>8 weeks) WT mice and 4.5-6.5 week-old KIC mice; both male and female animals were used. Group allocation (control or cKO) was based on Mendelian randomization. In other cases, mice were randomly assigned to treatment groups (αNetrin-1or Iso-antibody).

### Tamoxifen Injections

Tamoxifen (Sigma, Cat. #T5648-1G) dissolved in sunflower seed oil (Sigma, Cat. #S5007) was administered by intraperitoneal injection at a final dose of 100 mg/kg once daily for three consecutive days. A waiting period of 10 days was maintained between the last injection and either the sacrifice of the mice or the initiation of another treatment.

### Chronic pancreatitis induction

Mice were treated with cerulein (50 µg/kg; Sigma, Cat. #C9026) or phosphate-buffered saline (PBS; Thermo Fisher Scientific™, Cat. #16431755) (vehicle). Solutions were injected intraperitoneally every hour, 8 times a day, twice a week for 3.5 weeks. The mice were sacrificed 2 days after the final injection.

### Anti-Netrin-1 treatment

Anti-Netrin-1 monoclonal antibody (αNetrin-1) and IgG1 isotype control (Iso-antibody) were obtained from Netris Pharma (Lyon, France) (Supplementary Table 1). Antibodies diluted in PBS were administered via intraperitoneal injections twice weekly at a dose of 10 mg/kg. In the chronic pancreatitis model, each antibody injection was administered one day after the cerulein injection for a total duration of 3.5 weeks. In KIC mice, antibody injections were administered at 4.5 weeks of age and continued for 2 weeks. A detailed timeline is presented in the Figures.

### Tissue Harvest

The mice were anesthetized by intraperitoneal injections of 10 mg/kg xylazine (Rompun, Bayer) and 100 mg/kg ketamine (Imalgene, Merial). Intracardiac perfusion was then performed with 20 ml cold PBS, followed by 30 ml 4% paraformaldehyde (PFA; Electron Microscopy Sciences, Cat. #15714-S) in PBS. The pancreas, spleen, and surrounding nerve plexus were then removed as a single block and post-fixed by immersion in 4% PFA in PBS overnight at 4°C. The samples were rinsed three times with PBS. For RNAscope and immunostaining on cryosections, samples were incubated in 30% sucrose/1xPBS solution overnight at 4°C, embedded in OCT matrix (VWR, Cat. #361603E) and stored at –80°C. For wholemount immunostaining, samples were immediately processed.

### Immunofluorescence on cryostat sections

Immunofluorescence staining was performed on 20-µm thick cryostat sections. Tissue sections were washed with PBS and permeabilized in 0.1% Triton X-100 (Nacalai Tesque, Cat. #35501-15) in PBS for 15 min at RT. The sections were then blocked with gelatin blocking solution (0.02% gelatin; VWR Chemicals Cat. #24360.233, 0.25% Triton X-100 in PBS) for 1 h at RT and incubated with primary antibodies diluted in the blocking solution at 4°C overnight. The next day, sections were washed with 0.075% Triton X-100 in PBS and incubated with species-specific secondary antibodies for 2 h at RT. Sections were washed with 0.075% Triton X-100 in PBS and PBS, and incubated with NucBlue (Thermo Fisher, Cat. #R37606) for 6 min. Sections were then mounted in Aqua-Poly/Mount (Polysciences, Cat. #18606-20) and stored at 4°C until imaging. For anti-Netrin-1 and KI-67 immunostaining, antibody–antigen retrieval was performed prior to staining. To do so, sections were incubated at 85°C for 6 min in a freshly prepared 10mM citrate buffer pH 6 (sodium citrate dihydrate powder, Fisher Scientific, Cat. #BP327-500). In addition, Triton X-100 was excluded from all solutions for Netrin-1 staining. The primary and secondary antibodies used are listed in Supplementary Tables 2 and 3.

### RNAscope *in situ* hybridization

RNAscope *in situ* hybridization were performed on 14-µm thick cryostat sections using RNAscope Multiplex Fluorescent Reagent Kit v2 (Advanced Cell Diagnostics [ACD], Cat. #323100). RNA probes specific for Ntn1 (Cat. #407621-C2), Iba1 (Cat. #319141-C3), Dcc (Cat. #427491-C3), and Th (Cat. #317621) were previously designed and validated by ACD and are listed in Supplementary Table 4. Experiments were performed following the procedures in the RNAScope ISH manual from ACD (323100-USM), specifically, the protocol for fixed frozen sections using the RNAscope v2 kit (TSA). Sections were air-dried and pretreated with RNAscope hydrogen peroxide solution for 10 min at RT. Sections were washed with distilled water before boiling (98–102°C) in RNAscope 1x Target Retrieval solution for 6 min. Slides were immersed in distilled water for 15 s and then transferred for 3 min into 100% ethanol (VWR Chemicals, Cat. #20824.296) before air drying. After drying, a hydrophobic barrier was drawn around the tissue surface using the *ImmEdge* hydrophobic pen (Vector labs, Cat. #H-4000). On the next day, the slides were treated with RNAscope Protease

III reagent for 20 min at RT. The RNA Ntn1-C2 and Iba1-C3 probe combinations were prepared in the Probe Diluent supplied with the kit at the recommended concentration (1:50), and the Dcc-C3 probe was diluted directly in the Th-C1 probe (1:50). Slides were hybridized with the desired probe combination at 40°C in a HybEZ oven (ACD, Cat. #ACD_321720) for 2 h. Sections were washed with 1X RNA scope Wash Buffer for 2 min at RT after each of the following steps. Primary hybridization was amplified by incubating the sections with Z probe-specific preamplifiers and amplifiers in a step called FL V2 AmpX at 40°C in the HybEZ oven for 30 min, for each channel. The connection between the preamplifier and the amplifier bound to the Z probe was confirmed by adding Multiplex FL V2 HRP-CX at 40°C for 15 min. The detection step was completed using the following dyes: TSA Fluorescein (1:500; AKOYA Biosciences, Cat. #NEL701A001KT) for C1 probes, TSA Cyanine 3 (1:1500; AKOYA Biosciences, Cat. #NEL704A001KT) for C2 probes, and TSA Cyanine 5 (1:500; AKOYA Biosciences, Cat. #NEL705A001KT) for C3 probes prepared in 1X TSA buffer (provided in the kit) at 40°C in a HybEZ oven for 30 min. Finally, the hybridized structure on each mRNA was stabilized using the v2-HRP-blocker at 40°C for 15 min. Nuclei were stained with DAPI provided in the kit and sections were mounted with Aqua-Poly/Mount (Polysciences, Cat. #18606-20) and stored at 4°C until imaging.

### Whole-mount immunostaining and tissue clearing

The whole pancreas was immunostained and cleared using the iDISCO+ protocol ^46^. Samples were dehydrated with 20%, 40%, 60%, 80%, and 100% methanol solutions diluted in PBS (1 h each at RT), followed by bleaching with 5% hydrogen peroxide (Cat. #216763, Sigma– Aldrich) diluted in methanol, overnight at 4°C. The samples were then rehydrated with 100%, 80%, 60%, 40%, and 20% methanol solutions diluted in PBS (for 1 h each at RT), and permeabilized in 20% dimethyl sulfoxide (DMSO; Fisher Chemical, Cat. #D/4121/PB08), 0.16% Triton X-100 and 18.4 g/l glycine (VWR, Cat. #0167) in PBS at 37°C for 2 days. The samples were then incubated in blocking buffer: PTwH (0.2% Tween-20; VWR, Cat. #28829.183) and 0.01 mg/ml heparin (Sigma-Aldrich, Cat. #H7405) in PBS, 5% DMSO and 3% donkey serum (Deutscher, Cat. #S2170-100) at 37°C for 3 days. Primary antibodies diluted in blocking buffer were added to the sample and incubated at 37°C for 7 days. The samples were washed with PTwH before incubation with secondary antibodies in a blocking buffer at 37°C for 3 days. The antibodies used are listed in Supplementary Tables 2 and 3. In some cases, nuclear counterstaining was performed by incubation with TO-PRO^TM^-3 Iodide (642/661; Thermo Fisher Scientific, Cat. #T3605) diluted in DMSO (1:1000) for 1Ch 30Cmin at RT. After several washes with PTwH, the samples were dehydrated with 20%, 40%, 60%, 80%, and 100% methanol solutions diluted in PBS (1 h each at RT), and incubated overnight in 66% dichloromethane (DCM; Sigma-Aldrich, Cat. #270947) and 33% methanol at RT. Samples were delipidated by incubation in 100% DCM at RT, followed by immersion in dibenzyl ether (DBE; Sigma-Aldrich, Cat. #305197) for at least 3 h before imaging.

### Culture of sympathetic neurons

CSMG were isolated from adult S100b-GFP mice under a fluorescence dissection stereomicroscope based on the GFP expression of glial cells within the ganglion and collected in a cold Hank’s balanced salt solution (HBSS; Thermo Fisher Scientific, Cat. #14170112), 5 mM HEPES (Thermo Fisher Scientific, Cat. #15630080), 10 mM D-glucose (Thermo Fisher Scientific, Cat. #G/0500/60), and 100 U/ml penicillin–streptomycin (Thermo Fisher Scientific, Cat. #15140-122), pH 7.5. Tissues were incubated for 30 min at 37°C in HBSS, 5 mM CaCl_2_ (Thermo Fisher Scientific, Cat. #15641930), 2 mg/ml type II collagenase (Thermo Fisher Scientific, Cat. #17101-015) and 5 mg/ml dispase II (Thermo Fisher Scientific, Cat. #17105-041). The solution was refreshed, and incubation was continued for 30 min. The tissue was pelleted by centrifugation at 1300 × g for 10 min and washed with L-15 complete medium (Thermo Fisher Scientific, Cat. #11415049), 5% fetal calf serum (Gibco, Cat. #A5256801), and 100 U/ml penicillin–streptomycin. The cells were mechanically dissociated using fire-polished Pasteur pipettes by pipetting up and down 10–15 times. The cell suspension was applied to the top of a 12.5–28% Percoll gradient (Deutscher, Cat. #17-0891- 01) and centrifuged at 1300 × g for 20 min to separate the neurons from the debris. Neurons were resuspended in L-15 complete medium, centrifuged at 900 × g for 5 min, and resuspended in NB complete medium (Neurobasal A medium; Thermo Fisher Scientific, Cat. #10888022), B27 (Thermo Fisher Scientific, Cat. #12587010), 2 mM L-glutamine (Life tech invitro, Cat. #25030024), and 100 U/ml penicillin–streptomycin. Finally, the neurons were plated in 18-mm glass-bottomed dishes (Mattek, Cat. #P35G-1.5-20-C) coated with poly-D-lysine (1 mg/ml; Sigma–Aldrich, Cat. #P7280-5MG), and laminin (1 mg/ml; Sigma-Aldrich, Cat. #L2020-1MG) in NB complete medium supplemented with 50 ng/ml nerve growth factor (Alomone Laboratories, Cat. #N-240). Cells were cultured for 24 h at 37°C in 5% CO_2_.

### Immunostaining of neuronal culture

Cells were fixed with 4% PFA and 30% sucrose in PBS for 30 min at RT, washed in PBS, and permeabilized with 0.1% Triton X-100 in PBS for 15 min. Cells were blocked with 0.2% gelatin and 0.25% Triton X-100 in PBS for 1 h at RT and then incubated overnight with primary antibodies in blocking solution. After washing with PBS, the cells were incubated with secondary antibodies diluted in blocking solution for 2 h at RT. The antibodies used are listed in Supplementary Tables 2 and 3. Finally, nuclei were stained with Hoechst (1/5000) (Life Technologies, Cat. #R37605).

### Imaging

Immunohistochemistry (IHC) and RNAscope cryostat sections were imaged using the Axioimager M2 Apo1.2 apotome with 20x/0.8 (air) objective, and Zeiss LSM 880 airy scan confocal microscope with 20x/0.8 (air) objectives. Cleared pancreas were images using a light-sheet fluorescence microscope (UltraMicroscope Blaze, Miltenyi Biotec, Bergisch Gladbach, Germany) equipped with a 5.5 Megapixel sCMOS camera. Whole-pancreas imaging was done using a 1.1x/0.1 objective, whereas closer insets were imaged using a 4x/0.35 objective. The ImspectorPro 753 microscope control software version 7.3 was used to run the system.

Cultured CSMG neurons were imaged with an inverted AxioObserver Z1 Apo 2 using a 20x/0.8 objective.

### Data quantification

Data analysis was performed by investigators who were blinded to the genotype and anti-Netrin-1 antibody treatment. However, blinding to cerulein treatment was not feasible because its effects on ADM induction were apparent.

### 2D immunofluorescence images

Image analysis was performed using Fiji ^47^. The entire metaplastic or precancerous lesion was manually delineated as a region of interest (ROI) based on Sox9 or E-cadherin staining. In healthy or asymptomatic tissue, ROIs were defined using the autofluorescence of the pancreatic tissue.

#### Analysis of axon density

The NeuronJ plugin was used to trace and measure the total length of TH+, VAChT+ or CGRP+ axons within each ROI, based on the corresponding immunolabeling. The density of innervation was calculated by dividing the total axonal length by the area of the respective ROI.

#### Analysis of DCC expression by sympathetic axons

In each ROI, total TH+ axons and DCC+/TH+ axons were traced using NeuronJ plugin based on TH and DCC immunofluorescent signals. The corresponding axonal lengths were extracted, and the proportion of DCC-expressing sympathetic axons was calculated as the ratio of DCC+/TH+ axon length to total TH+ axon length.

#### Analysis of fluorescence signal density

Images thresholding was performed based on immunolabeling signal intensity to identify labeled regions. The area of these labeled regions was then measured and expressed as a proportion of the total ROI area.

#### Analysis of cell proliferation

For each imaged PanIN lesion, the number of epithelial cell and proliferating cell were counted using the multi-point tool, based on Sox9 and KI67 staining, respectively. The proportion of proliferating cell was calculated by dividing the number of KI67+/Sox9+ cells by the total number of Sox9+ cells.

#### Analysis of cell death

For each imaged PanIN lesion, the epithelial layer was manually delineated based on E-cadherin staining and the corresponding area was measured. The number of cells positive for activated caspase-3 was counted using the multi-point tool, and cell death density was calculated by dividing the number of counted cells by the measured area of the PanIN epithelium.

### 3D immunofluorescence images

3D reconstruction analysis was performed using Imaris x64 microscopy image analysis software 9.9.1 and 10.0.0 versions (Bitplane, Zurich, Switzerland). The stacked images were converted into Imaris files (.ims) using the Imaris File Converter. 3D reconstruction of the samples was performed using a 3D view in Imaris. For the analysis of sympathetic innervation, ROIs (healthy tissue or premalignant lesion) were selected based on tissue autofluorescence (imaged at 488 nm excitation) or CK19+ immunostaining. A mask was created around each ROI using the “Surface” tool and the ROI volume was collected in the statistics tab. In addition, for PanIN lesions, masks were also created around the lumens of epithelial lesions, and the volume of the lumen (empty space) was deduced from the ROI volume. Sympathetic axons in the selected ROIs were reconstructed either manually using the “Filament tracer” tool or automatically using the Imaris 10-AI-powered Filament tracer workflow. The following parameters were collected in the statistics tab: “Filament length (Sum)”—defined as the sum of the length of all axon branches, and “Filament no. dendrite branch points”—defined as the number of branch points in the entire axon network. The density of axons was normalized to the volume of the ROI (healthy or premalignant lesion) and was calculated as follows: *Dendrite length (Sum)* / (*Volume of the ROI* – *Volume of the lumens*)^1/3^. Axonal branching was calculated as: *Filament no. dendrite branch pts* x 1000 / *Dendrite length (Sum)*, and expressed as branch points/mm of the axon.

#### Analysis of RNAscope

Automated quantification of RNAscope data was performed using CellProfiler ^48^. To quantify the *Dcc* mRNA signal in the CSMG, sympathetic neuronal cell bodies were segmented based on the *Th* mRNA signal and the number. The number of *Dcc* dots per identified *Th+* cell was counted automatically based on a size range of 4–16 pixels (small: 4–9 pixels, medium: 9–12 pixels, large: 12–16 pixels). To quantify quantify the *Ntn1* mRNA signal in pancreatic tissue, ROI corresponding to asymptomatic tissue and premalignant lesion were first identified based on DAPI nuclear staining. Within each ROI, cell bodies were identified by the software as follow: nuclei were identified based on DAPI staining, filtered by size criteria (27–55 pixels), and annotated as primary objects. The nuclear object was dilated to a radius of 14 pixels to approximate the diameter of the entire cell body in the tissue. The reconstructed cell bodies were annotated and used to quantify the number of *Ntn1* dots per cell. *Ntn1* dots were quantified automatically with a size range of 3–10 pixels (small dots: 3–5 pixels, medium dots: 5–7 pixels, large dots: 7–10 pixels). The average level of transcript expression was calculated based on the signal intensity of each type of dot and the assumption that small dots corresponded to a single mRNA: number of small dots + (number of medium dots x 2) + (number of large dots x 3).

#### Analysis of 2D primary cultures

Fluorescence images of the cultured CSMG neurons were automatically analyzed based on TH staining using the open-source MATLAB-based software developed by ^49^ to determine the ‘total neurite length’ for each neuron. The *in vitro* culture experiment was repeated at least three times and consistent data replication was observed each time.

### sNuc-seq data analysis

Chronic Pancreatitis sNuc-Seq data from two donors was obtained from the Human Cell Atlas Data Portal (https://explore.data.humancellatlas.org/projects/b3938158-4e8d-4fdb-9e13-9e94270dde16). Data were analyzed using Scanpy on the Galaxy Europe server (https://singlecell.usegalaxy.eu/)^50^. Low-quality nuclei identified by a low (<200) or high (>5000) gene counts and nuclei containing more than 3% of mitochondrial reads were excluded from downstream analyses. Counts were normalized, fitting total counts to 10,000 per cell, log-transformed, and the most variable genes were identified as those with a normalized dispersion amount higher than 0.5. Data were scaled to unit variance and zero mean, and the dimensionality of the data was reduced by principal component analysis (PCA) (30 components). k-nearest neighbors (kNN) with k=15 was used to compute a neighboring graph, and clustering was performed using the Louvain algorithm (resolution = 0.5) and visualized with UMAP. Marker genes were identified using the Wilcoxon rank sum test and clusters were manually assigned to known cell types based on established marker genes^22^.

### Human tissues

Formalin-fixed, paraffin-embedded (FFPE) human PDAC tissues were obtained from the Biobank Shared Resource (BSR) at the Wilmot Cancer Institute. The BSR processes, stores, and disburses biospecimens and subject data from consenting subjects undergoing a cancer screening or treatment at the University of Rochester Medical Center. All biobanking and clinical research activities performed by the BSR were developed in accordance with, and strictly adhere to, National Cancer Institute (NCI BBRB) and International Society for Biological and Environmental Repositories (ISBER) best practices. Paraffin blocks were sectioned at 4 µm and dried for 1 hour at 65C°C. Sections were rehydrated and subjected to antigen retrieval (36 minutes at 95C°C in Tris/EDTA CC1 buffer; Roche, Cat. #950-500), followed by blocking of endogenous peroxidases using Inhibitor CM (Roche, Cat. #760- 4840). Immunohistochemistry was performed using an automated immunostainer (Ventana Discovery XT, Roche) with the Discovery anti-rabbit HQ HRP kit and the ChromoMap DAB Kit (Roche, Cat. #760-159). Sections were incubated with a primary antibody against Netrin- 1, followed by detection with anti-rabbit HQ and amplification with anti-HQ horseradish peroxidase (Roche). Signal was visualized using 3,3′-diaminobenzidine as the chromogenic substrate and counterstained with Gill’s haematoxylin. Finally, sections were scanned at 20X magnification using the Pannoramic Scan II system (3D Histech, Budapest, Hungary). Antibody details are provided in Supplementary Tables 2 and 3.

### Statistical analysis

In the absence of preliminary data, the sample sizes were determined using the resource equation. After the pilot experiments provided information on the effect size, the sample sizes were calculated by power analysis using the BiostaTGV calculator (https://biostatgv.sentiweb.fr/). The statistical power was set at 80% with a significance level of 5%. GraphPad Prism version 9 (GraphPad Software Inc.) was used for the statistical analysis. All analyzed data were tested for normality using the Kolmogorov–Smirnov, D’Agostino–Pearson omnibus, or Shapiro–Wilk tests. Based on the normality results and the size of the manipulated data, the statistical tests performed included the two-tailed t-test and Mann–Whitney test (all performed as two-tailed tests). To compare three or more different datasets, one-way ANOVA (multiple comparisons) was used, and the Kruskal–Wallis test was performed for non-parametric multiple comparisons. The frequencies between the two groups were compared using the Chi-squared test. The Figure legends provide detailed information on the statistical tests and number of animals used for each experiment.

## Data availability

The data that support the findings of this study are available within the paper and its supplementary information files. Materials and full microscopy image data sets are available from the corresponding author upon request.

## Acknowledgments

We thank all members of our laboratory for their insightful contributions throughout this study, and Sophie Chauvet and Pascale Durbec for their critical reading of the manuscript. We are grateful to Martha Montserrat Rangel Sosa for assistance with the chronic pancreatitis model, Antoine Amabile for RNAscope data analysis, and the staff of the IBDM animal facility for their support. We thank the imaging facility at IBDM, member of the National Infrastructure France-BioImaging (https://ror.org/01y7vt929) supported by the French National Research Agency (ANR-24-INBS-0005 FBI BIOGEN). We also thank Rejane Rua for providing the LysM-Cre mice, Aziz Moqrich and David Ginty for the Th-CreERT2 mice, and Tobby Lawrence and Marc Bajenoff for the anti-CD206 and anti-CD163 antibodies. Human tissues were obtained from the Biobank Shared Resource at the Wilmot Cancer Institute, supported in part by the University of Rochester Wilmot Cancer Institute Support Grant #P30CA272302. The content is solely the responsibility of the authors and does not necessarily represent the official views of the National Institutes of Health. Some calculations were performed using the Galaxy server, partially funded by the German Federal Ministry of Education and Research (BMBF grant 031 A538A de.NBI-RBC) and the Ministry of Science, Research and the Arts Baden-Württemberg (MWK), within the framework of LIBIS/de.NBI Freiburg. This work was supported by the Centre National de la Recherche Scientifique (CNRS), Aix Marseille Université (AMU), and the Institut National de la Santé et de la Recherche Médicale (INSERM), France, as well as grants from the Fondation pour la Recherche Médicale (EQ202103012957), INCa, the Ligue contre le cancer, and Fondation ARC (PAIR Pancreas 186738) to FM. HH and KS received fellowships from the French Ministry of Higher Education, Research, and Innovation (MESRI).

## Author contributions

F.M. conceived the study and wrote the initial draft of the manuscript. H.H. and A.B. designed and carried out the experiments, including data collection and analysis, and co-wrote the manuscript. K.S. conducted and analyzed the in vitro culture experiments. M.H. contributed to the 3D image analyses. D.C., N.R., and N.G. provided and processed human tissue samples. P.M. supplied reagents and experimental animals. All authors reviewed and contributed to the final revision of the manuscript.

## Competing interests

P.M. is a shareholder of Netris Pharma.

The other authors declare no competing interests.

**Materials & Correspondence:** Fanny Mann.

## References

1. Mancusi, R. & Monje, M. The neuroscience of cancer. Nature 618, 467–479 (2023).

2. Ayala, G. et al. Cancer-Related Axonogenesis and Neurogenesis in Prostate Cancer. Clin Cancer Res 14, 7593–7603 (2009).

3. Magnon, C. et al. Autonomic nerve development contributes to prostate cancer progression. Science 341, 1236361 (2013).

4. Stopczynski, R. E. et al. Neuroplastic Changes Occur Early in the Development of Pancreatic Ductal Adenocarcinoma. Cancer Res 74, 1718–1727 (2014).

5. Sinha, S. et al. PanIN Neuroendocrine Cells Promote Tumorigenesis via Neuronal Cross-talk. Cancer Res 77, 1868–1879 (2017).

6. Guillot, J. et al. Sympathetic axonal sprouting induces changes in macrophage populations and protects against pancreatic cancer. Nat Commun 13, 1985 (2022).

7. Song, Y. et al. Enriching the Housing Environment for Mice Enhances Their NK Cell Antitumor Immunity via Sympathetic Nerve–Dependent Regulation of NKG2D and CCR5. Cancer Res 77, 1611–1622 (2017).

8. Stoeckli, E. T. Understanding axon guidance: are we nearly there yet? Development 145, dev151415 (2018).

9. Biankin, A. V et al. Pancreatic cancer genomes reveal aberrations in axon guidance pathway genes. Nature 491, 399–405 (2012).

10. Murphy, S. J. et al. Genetic Alterations Associated With Progression From Pancreatic Intraepithelial Neoplasia to Invasive Pancreatic Tumor. Gastroenterology 145, 1098–1109.e1 (2013).

11. Chédotal, A., Kerjan, G. & Moreau-Fauvarque, C. The brain within the tumor: new roles for axon guidance molecules in cancers. Cell Death Differ 12, 1044–1056 (2005).

12. Harburg, G. C. & Hinck, L. Navigating breast cancer: axon guidance molecules as breast cancer tumor suppressors and oncogenes. J Mammary Gland Biol Neoplasia 16, 257–270 (2011).

13. Brisset, M., Grandin, M., Bernet, A., Mehlen, P. & Hollande, F. Dependence receptors: new targets for cancer therapy. EMBO Mol Med 13, e14495 (2021).

14. Prévot, P.-P. et al. Role of the ductal transcription factors HNF6 and Sox9 in pancreatic acinar-to-ductal metaplasia. Gut 61, 1723–1732 (2012).

15. Liou, G.-Y. et al. Macrophage-secreted cytokines drive pancreatic acinar-to-ductal metaplasia through NF-κB and MMPs. Journal of Cell Biology 202, 563–577 (2013).

16. van Gils, J. M. et al. The neuroimmune guidance cue netrin-1 promotes atherosclerosis by inhibiting macrophage emigration from plaques. Nat Immunol 13, 136–143 (2012).

17. Finci, L. I. et al. The crystal structure of netrin-1 in complex with DCC reveals the bifunctionality of netrin-1 as a guidance cue. Neuron 83, 839–849 (2014).

18. Grandin, M. et al. Structural Decoding of the Netrin-1/UNC5 Interaction and its Therapeutical Implications in Cancers. Cancer Cell 29, 173–185 (2016).

19. Meijers, R., Smock, R. G., Zhang, Y. & Wang, J.-H. Netrin Synergizes Signaling and Adhesion through DCC. Trends Biochem Sci 45, 6–12 (2020).

20. Hadi, T. et al. Macrophage-derived netrin-1 promotes abdominal aortic aneurysm formation by activating MMP3 in vascular smooth muscle cells. Nat Commun 9, 5022 (2018).

21. Sharma, M. et al. Netrin-1 Alters Adipose Tissue Macrophage Fate and Function in Obesity. Immunometabolism 1, e190010 (2019).

22. Tosti, L. et al. Single-Nucleus and In Situ RNA-Sequencing Reveal Cell Topographies in the Human Pancreas. Gastroenterology 160, 1330–1344.e11 (2021).

23. Brunet, I. et al. Netrin-1 controls sympathetic arterial innervation. J Clin Invest 124, 3230– 3240 (2014).

24. Yebra, M. et al. Recognition of the neural chemoattractant Netrin-1 by integrins alpha6beta4 and alpha3beta1 regulates epithelial cell adhesion and migration. Dev Cell 5, 695–707 (2003).

25. Gao, R. et al. Macrophage-derived netrin-1 drives adrenergic nerve–associated lung fibrosis. J Clin Invest 131, (2021).

26. Castets, M. & Mehlen, P. Netrin-1 role in angiogenesis: to be or not to be a pro-angiogenic factor? Cell Cycle 9, 1466–1471 (2010).

27. Dudgeon, C. et al. Netrin-1 feedforward mechanism promotes pancreatic cancer liver metastasis via hepatic stellate cell activation, retinoid, and ELF3 signaling. Cell Rep 42, 113369 (2023).

28. Dumartin, L. et al. Netrin-1 Mediates Early Events in Pancreatic Adenocarcinoma Progression, Acting on Tumor and Endothelial Cells. Gastroenterology 138, 1595–1606.e8 (2010).

29. Lei, B., Schwinn, D. A. & Morris, D. P. Stimulation of α1a Adrenergic Receptors Induces Cellular Proliferation or Antiproliferative Hypertrophy Dependent Solely on Agonist Concentration. PLoS One 8, e72430 (2013).

30. Wang, W., Guo, X. & Dan, H. α2A-Adrenergic Receptor Inhibits the Progression of Cervical Cancer Through Blocking PI3K/AKT/mTOR Pathway. Onco Targets Ther 13, 10535–10546 (2020).

31. Wu, C. S., Tsao, D. A. & Chang, H. R. Beta2-adrenergic receptor agonist inhibits keratinocyte proliferation by mechanisms involving nitric oxide. Postepy Dermatol Alergol. 38, 396 (2020).

32. Renz, B. W. et al. β2 adrenergic-neurotrophin feed-forward loop promotes pancreatic cancer. Cancer Cell 33, 75–90.e7 (2018).

33. Thiel, V. et al. Characterization of single neurons reprogrammed by pancreatic cancer. Nature (2025) doi:10.1038/S41586-025-08735-3.

34. Cassier, P. A. et al. Netrin-1 blockade inhibits tumour growth and EMT features in endometrial cancer. Nature 620, 409–416 (2023).

35. Braxton, A. M. et al. 3D genomic mapping reveals multifocality of human pancreatic precancers. Nature 629, 679–687 (2024).

36. Kryza, D. et al. From netrin-1-targeted SPECT/CT to internal radiotherapy for management of advanced solid tumors. EMBO Mol Med 15, e16732 (2023).

37. Dominici, C. et al. Floor-plate-derived netrin-1 is dispensable for commissural axon guidance. Nature 545, 350–354 (2017).

38. Clausen, B. E., Burkhardt, C., Reith, W., Renkawitz, R. & Förster, I. Conditional gene targeting in macrophages and granulocytes using LysMcre mice. Transgenic Res 8, 265–277 (1999).

39. Gu, G., Dubauskaite, J. & Melton, D. A. Direct evidence for the pancreatic lineage: NGN3+ cells are islet progenitors and are distinct from duct progenitors. Development 129, 2447– 2457 (2002).

40. Abraira, V. E. et al. The Cellular and Synaptic Architecture of the Mechanosensory Dorsal Horn. Cell 168, 295–310.e19 (2017).

41. Krimpenfort, P. et al. Deleted in colorectal carcinoma suppresses metastasis in p53-deficient mammary tumours. Nature 482, 538–541 (2012).

42. Srinivas, S. et al. Cre reporter strains produced by targeted insertion of EYFP and ECFP into the ROSA26 locus. BMC Dev Biol 1, 4 (2001).

43. Vives, V., Alonso, G., Solal, A. C., Joubert, D. & Legraverend, C. Visualization of S100B- positive neurons and glia in the central nervous system of EGFP transgenic mice. J Comp Neurol 457, 404–419 (2003).

44. Jackson, E. L. et al. Analysis of lung tumor initiation and progression using conditional expression of oncogenic K-ras. Genes Dev 15, 3243–3248 (2001).

45. Aguirre, A. J. et al. Activated Kras and Ink4a/Arf deficiency cooperate to produce metastatic pancreatic ductal adenocarcinoma. Genes Dev 17, 3112–3126 (2003).

46. Renier, N. et al. Mapping of Brain Activity by Automated Volume Analysis of Immediate Early Genes. Cell 165, 1789–1802 (2016).

47. Schindelin, J., et al. Fiji: an open-source platform for biological-image analysis. Nat Methods 9, 676–682 (2012).

48. Carpenter, A. E. et al. CellProfiler: image analysis software for identifying and quantifying cell phenotypes. Genome Biol 7, R100 (2006).

49. Zehtabian, A., Fuchs, J., Eickholt, B. J. & Ewers, H. Automated Analysis of Neuronal Morphology through an Unsupervised Classification Model of Neurites. Preprint at 10.1101/2022.03.01.482454 (2022).

50. Abueg, L. A. L. et al. The Galaxy platform for accessible, reproducible, and collaborative data analyses: 2024 update. Nucleic Acids Res 52, W83–W94 (2024).

