## Supplementary information for "Neural function of Netrin-1 in precancerous lesions of the pancreas"

### Supplementary Figures

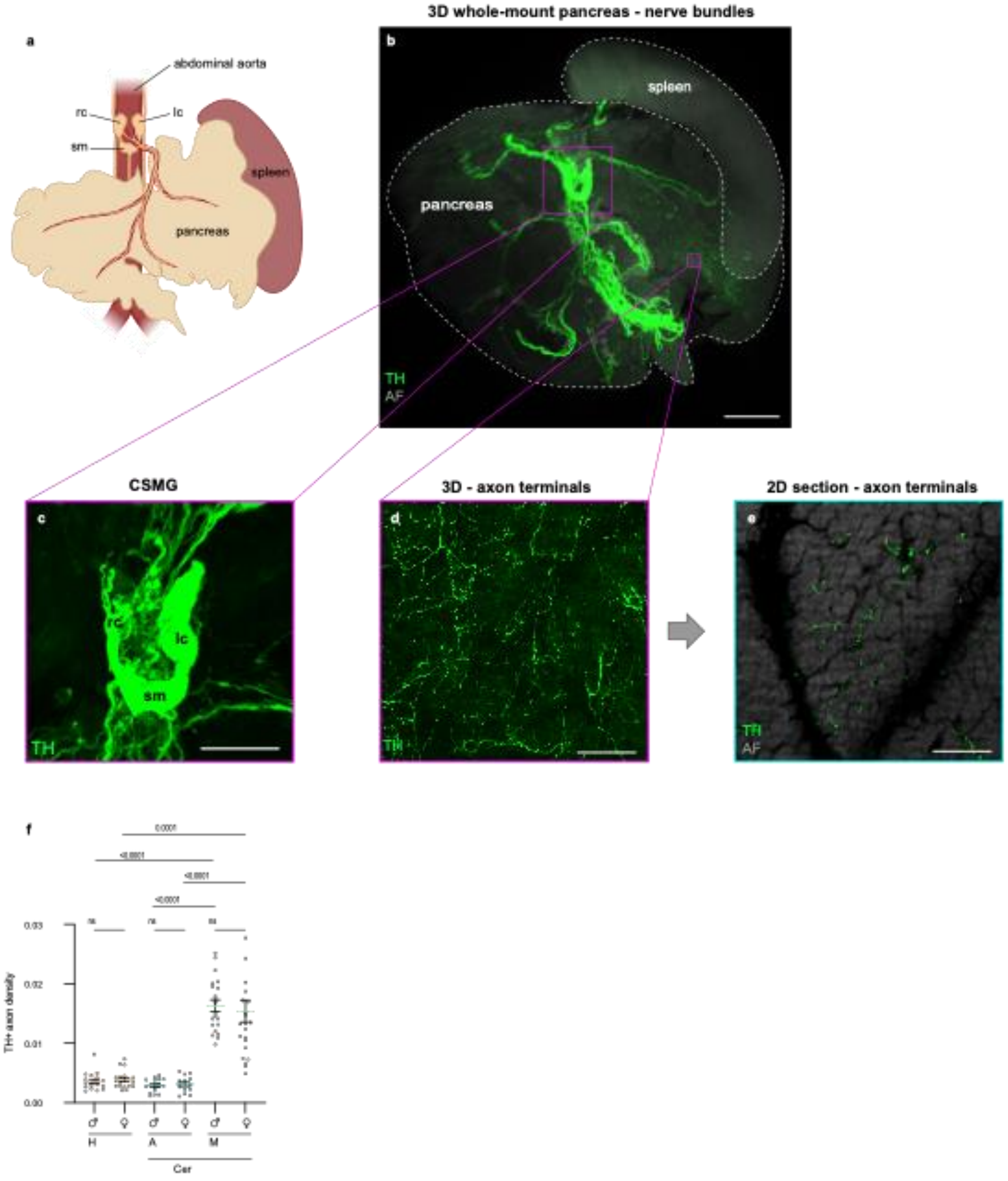


**Supplementary Fig. S1**: **Sympathetic innervation of the pancreas and its remodeling in metaplastic pancreatic lesions.**

**a**, Schematic showing the innervation of the mouse pancreas by projections from sympathetic neurons of the celiac-superior mesenteric ganglia. rc, right celiac ganglion; lc, left celiac ganglion; sm, superior mesenteric ganglion.

**b-c,** Maximum intensity projection of 3D image stacks of an 8-week-old mouse pancreas showing the TH+ sympathetic nerve bundles arising from the coeliac-superior mesenteric ganglion complex (CSMG). Autofluorescence (AF) allows the visualization of the entire pancreas and the spleen, which was left attached to the pancreas during dissection. The CSMG (c) consists of distinct ganglionic subunits (rc., right celiac ganglion; lc, left celiac ganglion; sm, superior mesenteric ganglion). After entering the pancreas, $TH^+$ sympathetic nerve bundles branch out to innervate the pancreas. Scale bars: 2000 µm (a), and 700 µm (b).

**d-e,** Maximum intensity projection of 3D image stacks (d) and 2D optical section (e) of the same region of the exocrine pancreas of an 8-week-old mouse pancreas showing TH+ sympathetic axon terminals. Scale bars: 100 µm.

**f**, Data on the $TH^+$ sympathetic axon density in the exocrine pancreas of healthy mice (H), metaplastic lesions (M), and adjacent exocrine tissue (A) of cerulein-treated (Cer) mice, shown in Fig. 1e, were separated by sex. Data are expressed as mean ± SEM. $n$ = 18 ROIs from 3 healthy males, 21 metaplastic, and 17 adjacent ROIs from 3 cerulein-treated males. $n$ = 18 ROIs from 3 healthy females, 23 metaplastic, and 14 adjacent ROIs from 3 cerulein-treated females. Significant p values are indicated on the graph (ns: $p > 0.05$) (Kruskal–Wallis test).

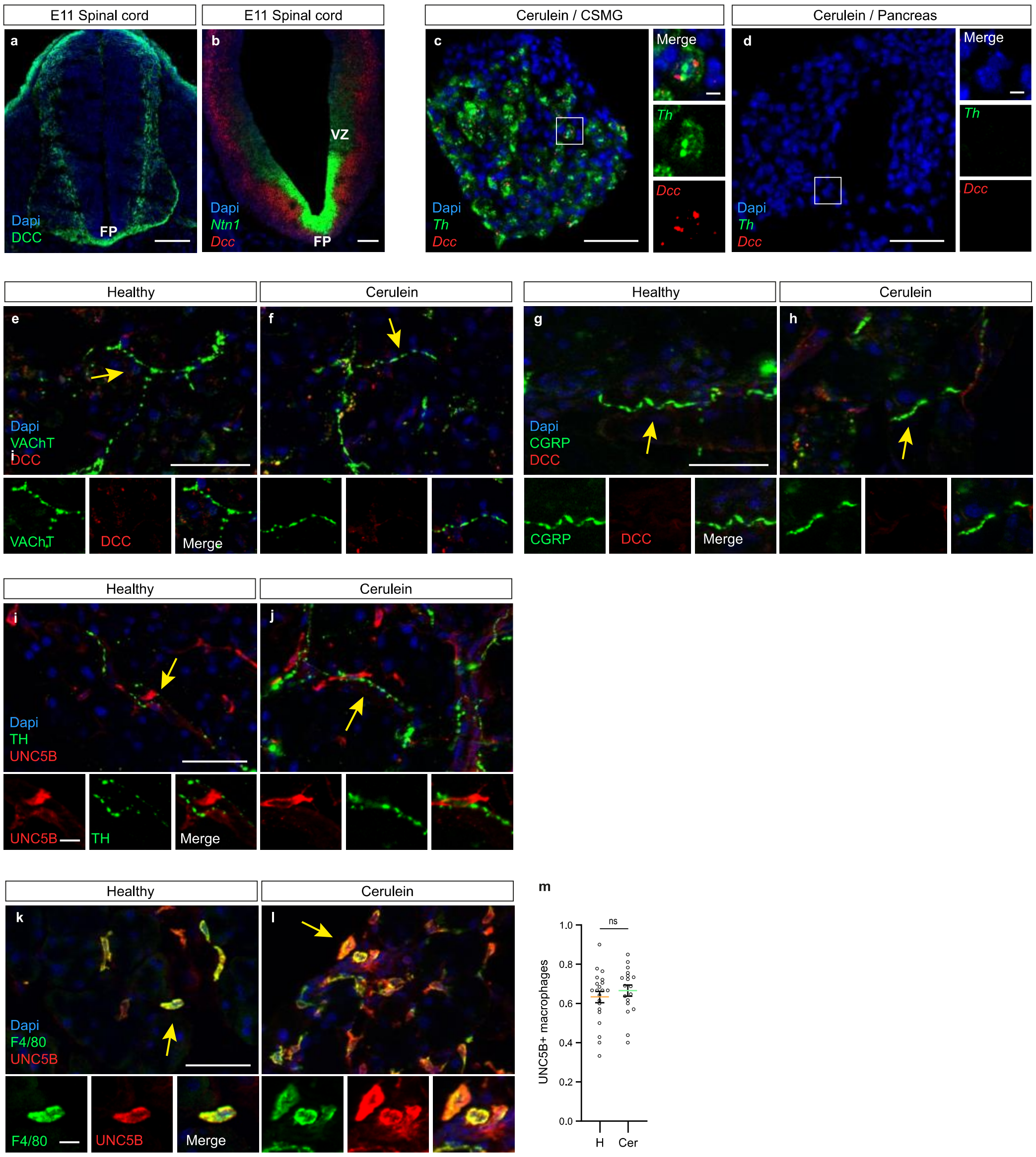


**Supplementary Fig. S2**: **DCC and UNC5B receptor expression in healthy and metaplastic pancreas.**

**a,** Validation of the DCC antibody for immunohistochemical (IHC) staining of mouse embryonic (E)11 spinal cord sections. The DCC labels the projections of dorsal commissural neurons to the ventral midline floor plate (FP). Scale bar: 100 µm.

**b,** Validation of the RNAscope probes for *Dcc* and *Ntn-1*. The probes were hybridized to E11 spinal cord sections. The *Ntn-1* probe typically binds to floor plate cells and ventral two-thirds of the ventricular zone (VZ), whereas the Dcc probe binds to a broad lateral domain. Scale bar: 100 µm.

**c-d,** Representative RNAscope *in situ* hybridization images of *Th* mRNA and *Dcc* mRNA in a tissue section from a cerulein-treated mouse containing both the celiac superior mesenteric ganIa (CSMG) (c) and metaplastic pancreas (d). Although *Dcc* was expressed in sympathetic CSMG neurons, no signal was detected in the corresponding metaplastic pancreas. DAPI was used for nuclear staining. These images represent the results from three mice analyzed. Scale bar: 50 µm and 5 µm (insets)

**e-f,** Representative images of IHC staining for VAChT (a marker of cholinergic neurons) and DCC in sections of the healthy exocrine pancreas (e) and cerulein-induced metaplastic pancreatic lesions (f). DAPI was used for nuclear staining. These images represent results from three mice analyzed per condition, in which zero VaCht+ axons with detectable DCC expression were found. Scale bar: 50 µm.

**g-h**, Representative images of IHC staining for CGRP (a marker of peptidergic sensory neurons) and DCC in sections of healthy exocrine pancreas (g) and cerulein-induced metaplastic pancreatic lesions (h). DAPI was used for nuclear staining. These images represent results from three mice analyzed per condition, in which zero CGRP+ axons with detectable DCC expression were found. Scale bar: 50 µm.

**i-j,** Representative images of IHC staining for TH and UNC5B in sections of healthy exocrine pancreas (i) and cerulein-induced metaplastic pancreatic lesions (j). DAPI was used for nuclear staining. These images represent results from three mice analyzed per condition, in which zero CGRP+ axons with detectable DCC expression were found. Scale bar: 50 µm and 10 µm (insets).

**k-l,** Representative images of IHC staining for F4/80 (a macrophage marker) and UNC5B in sections of healthy (k) and cerulein-treated (l) pancreas. DAPI was used for nuclear staining. These images represent results from three mice analyzed per condition. Scale bar: 50 µm and 10 µm insets.

**m**, Quantification of the proportion of F4/80+ macrophages expressing UNC5B in healthy (H) and cerulein-treated (Cer) pancreas. Data are presented as mean ± SEM. $n = 21$ healthy and 18 metaplastic ROIs from 3 mice/group. ns: $p > 0.05$ (Mann–Whitney test).

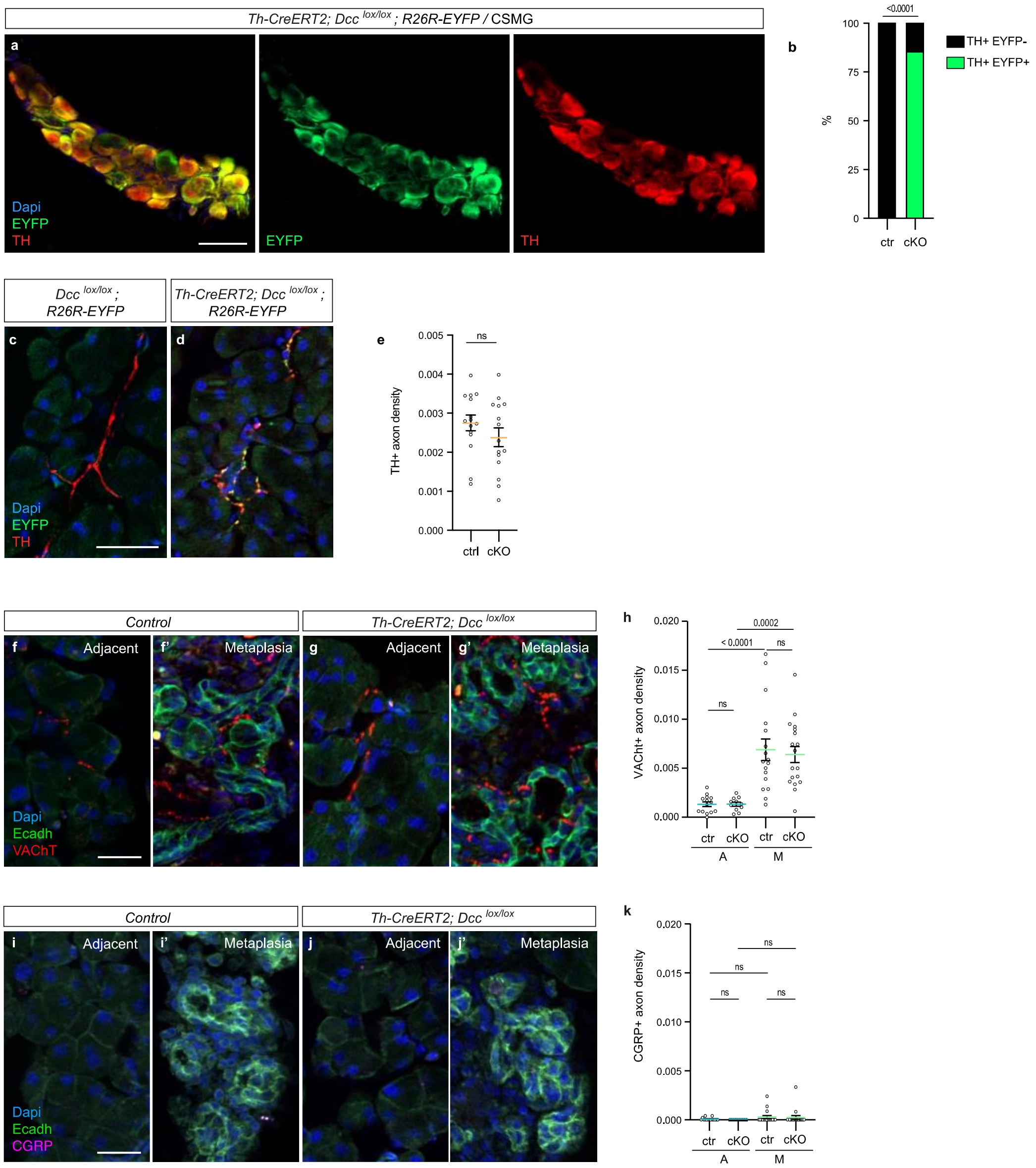


**Supplementary Fig. S3**: **Efficiency and impact of *Dcc* deletion in sympathetic neurons.**

**a**, Representative images of immunohistochemical (IHC) staining for TH and EYFP in sections through the celiac superior mesenteric ganglia (CSMG) of *Th-CreERT2–Dcc*$^{lox/lox}$*;* and R26R-EYFP mice 10 days after receiving three injections of tamoxifen. DAPI was used for nuclear staining. Scale bar: 50 µm.

**b**, Percentage of TH+ sympathetic neurons showing tamoxifen-dependent recombination of the R26R-EYFP allele (TH$^{+}$/EYFP$^{+}$ neurons) in the CSMG of *Th-CreERT2–Dcc*$^{lox/lox}$*;* R26R-EYFP mice (cKO) and control Cre-negative *Dcc*$^{lox/lox}$; R26R-EYFP (ctr) mice. $n$ = 136 TH+ neurons from 3 ctr mice and 88 TH+ neurons from 3 cKO mice. p-values are indicated on the graph (Chi-squared test).

**c-d,** Representative images of IHC staining for TH and EYFP in the exocrine pancreas of tamoxifen-treated *Th-CreERT2–Dcc*$^{lox/lox}$*;* R26R-EYFP mice (d) and control Cre-negative *Dcc*$^{lox/lol}$R26R-EYFP mice (c). Scale bar: 50 µm.

**e,** Quantification of TH$^{+}$ sympathetic axon density in the exocrine pancreas of tamoxifen-treated *Th-CreERT2–Dcc*$^{lox/lox}$*;* R26R-EYFP (cKO) and *Dcc*$^{lox/lox}$; R26R-EYFP (ctr) mice. Data are presented as mean ± SEM. $n = 15$ sections from 3 mice per genotype. ns: $p > 0.05$ (Mann–Whitney test).

**f-g,** Representative images of IHC staining for VAChT and E-cadherin (Ecadh) in metaplastic pancreatic lesions (f' and g') and adjacent exocrine tissue (f and g) of cerulein-treated control and *Th-CreERT2–Dcc* $^{lox/lox}$ mice. DAPI was used for nuclear staining. Scale bar: 50 µm.

**h**, Quantification of VAChT$^{+}$ axon density in metaplastic pancreatic lesions (M) and adjacent tissues (A) of cerulein-treated control (ctr) and *Th-CreERT2–Dcc* $^{lox/lox}$ (cKO) mice. Data are presented as mean ± SEM. $n = 17$ metaplastic and 14 adjacent ROIs from 3 control mice; 18 metaplastic and 11 adjacent ROIs from 3 cKO mice. ns: $p > 0.05$ (Kruskal–Wallis test).

**i-j,** Representative images of IHC staining for CGRP and E-cadherin (Ecadh) in metaplastic pancreatic lesions (i' and j') and adjacent exocrine tissue (i and j) of cerulein-treated control and *Th-CreERT2–Dcc* $^{lox/lox}$ mice. DAPI was used for nuclear staining. Scale bar: 50 µm.

**k**, Quantification of CGRP$^{+}$ axon density in metaplastic pancreatic lesions (M) and adjacent tissues (A) of cerulein-treated control (ctr) and *Th-CreERT2–Dcc* $^{lox/lox}$ (cKO) mice. Data are presented as mean ± SEM. $n = 17$ metaplastic and 15 adjacent ROIs from 3 control mice; 18 metaplastic and 10 adjacent ROIs from 3 cKO mice. ns: $p > 0.05$ (Kruskal–Wallis test).

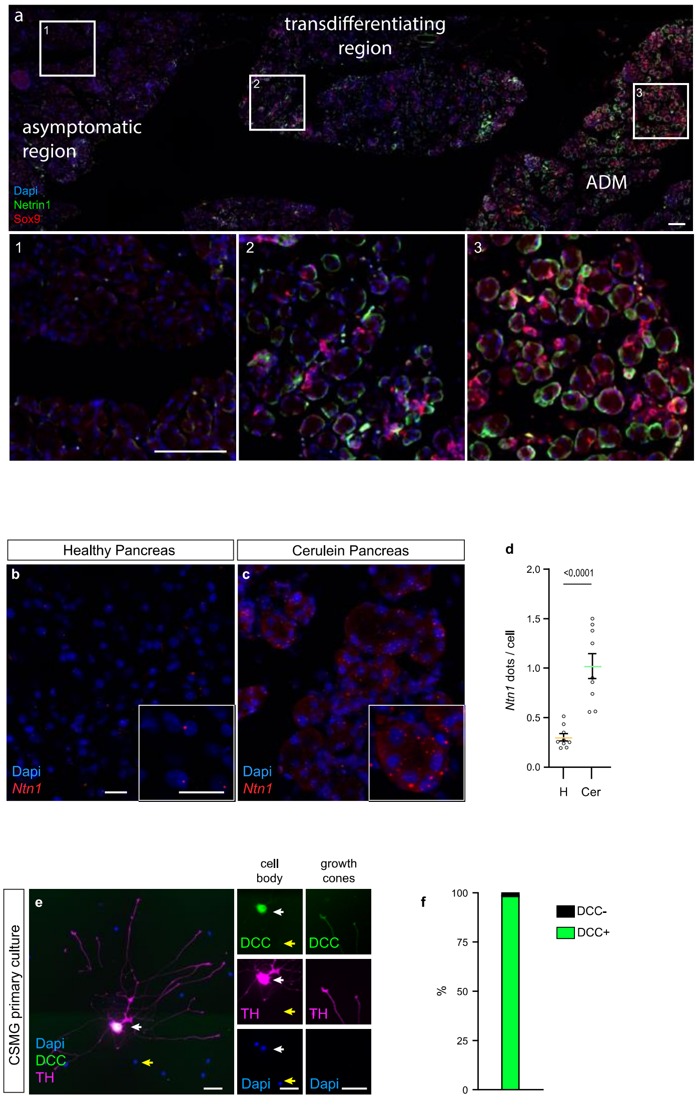


**Supplementary Fig. S4: Netrin-1 and DCC expression in cerulein-treated pancreas and cultured sympathatic neurons.**

**a**, Representative images of immunohistochemical staining for Netrin-1 and Sox9 in a section of cerulein-treated pancreas. Netrin-1 signal is high in regions with Sox9+ Acinar to Ductal Metaplasia (ADM) (3), sparse in the transition zone where Sox9+ ductal markers begin to be expressed (2), and absent in the asymptomatic region (1). DAPI was used for nuclear staining. Scale bar: 100 µm.

**b-c**, Representative images of RNAscope *in situ* hybridization of *Ntn1* mRNA in sections of the healthy exocrine pancreas (b) and cerulein-induced metaplastic pancreatic lesions (c). DAPI was used for nuclear staining. Scale bar: 20 µm.

**d**, Quantification of *Ntn1* mRNA (dots) in CSMG sympathetic neurons from healthy (H) and cerulein-treated (Cer) mice. Data are presented as mean ± SEM. $n = 9$ sections from 3 healthy mice and 8 sections from 3 cerulein-treated mice. p values are indicated in the graph (unpaired t-test).

**e,** Immunolabeling of TH and DCC in cultured dissociated cells from the celiac superior mesenteric ganglia (CSMG). DCC is expressed on the cell bodies and axonal growth cones of $TH^{+}$ neurons (white arrow), but not on $TH^{-}$ satellite glial cells (yellow arrow). DAPI was used for nuclear staining. Scale bar: 50 μm.

**f**, Quantification of DCC expression in cultured CSMG neurons: 98.3% (357 out of 363) TH+ neurons expressed DCC. Data are from three independent cultures (n=3).

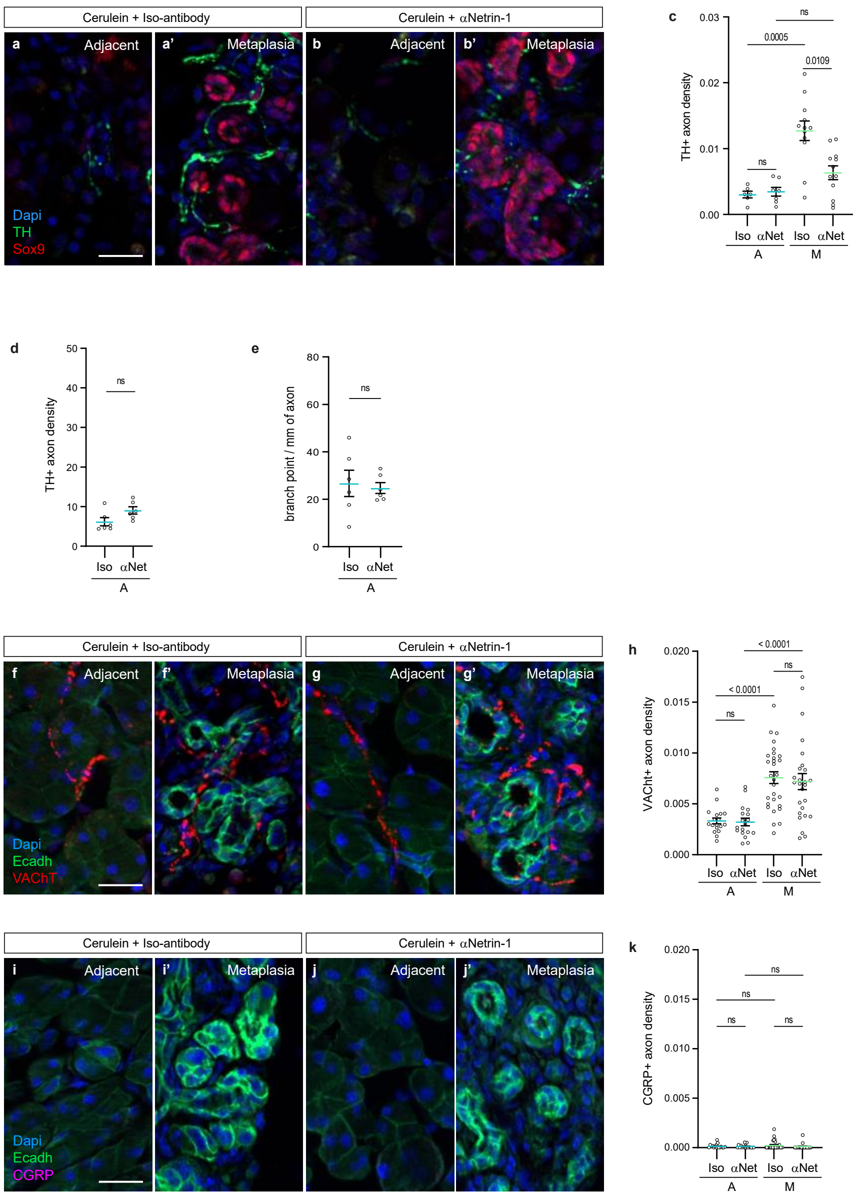


**Supplementary Fig. S5: Impact of Netrin-1 inhibition on metaplastic innervation.**

**a-b**, Representative images of IHC staining for TH and Sox9 in sections of cerulein-induced metaplastic pancreatic lesions (a' and b') and adjacent tissue (a and b) from mice treated with anti-Netrin-1 antibody (αNetrin-1) or isotype control antibody (Iso-antibody). DAPI was used for nuclear staining. Scale bar: 25 µm.

**c**, Quantification of $TH^{+}$ sympathetic axon density in cerulein-induced metaplastic pancreatic lesions (M) and adjacent tissues (A) from mice treated with Iso-antibody (Iso) or αNetrin-1 (αNet). Data are presented as mean ± SEM. $n = 12$ metaplastic and 6 adjacent ROIs (Iso) and 12 metaplastic and 4 adjacent ROIs (αNet) from 4 mice/group. Significant p values are indicated on the graph (ns: $p > 0.05$) (Kruskal–Wallis test).

**d-e,** 3D quantification of $TH^+$ sympathetic axon density (d) and axon branching (e) in tissues adjacent to metaplastic lesions (A) in mice treated with Iso-antibody (Iso) or αNetrin-1 (αNet) antibodies. Data are presented as mean ± SEM. $n = 6$ ROIs (Iso) and 6 ROIs (αNet) from 3 mice per group. ns: $p > 0.05$ (Mann-Whitney).

**f-g,** Representative images of IHC staining for VAChT and E-cadherin (Ecadh) in sections of cerulein-induced metaplastic pancreatic lesions (f' and g') and adjacent tissue (f and g) from mice treated with Iso-antibody or αNetrin-1. DAPI was used for nuclear staining. Scale bar: 25 µm.

**h**, Quantification of $VAChT^+$ axon density in cerulein-induced metaplastic pancreatic lesions (M) and adjacent tissues (A) from mice treated with Iso-antibody (Iso) or αNetrin-1 (αNet). Data are presented as mean ± SEM. $n = 29$ metaplastic and 18 adjacent ROIs (Iso) and 26 metaplastic and 18 adjacent ROIs (αNet) from 3 mice/group. ns: $p > 0.05$ (Kruskal–Wallis test).

**i-j,** Representative images of IHC staining for CGRP and E-cadherin (Ecadh) in sections of cerulein-induced metaplastic pancreatic lesions (i' and j') and adjacent tissue (i and j) from mice treated with Iso-antibody or αNetrin-1. DAPI was used for nuclear staining. Scale bar: 25 µm.

**k**, Quantification of $CGRP^+$ axon density in cerulein-induced metaplastic pancreatic lesions (M) and adjacent tissues (A) from mice treated with Iso-antibody (Iso) or αNetrin-1 (αNet). Data are presented as mean ± SEM. $n = 29$ metaplastic and 17 adjacent ROIs (Iso) and 26 metaplastic and 18 adjacent ROIs (αNet) from 3 mice/group. ns: $p > 0.05$ (Kruskal–Wallis test).

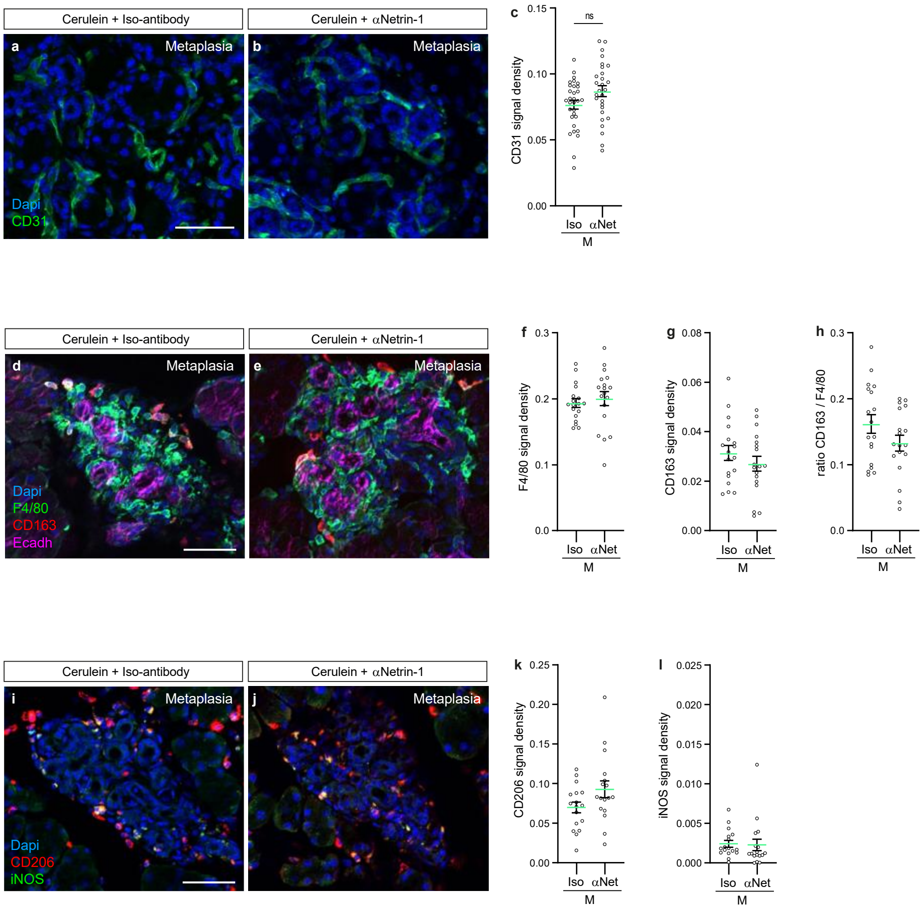


**Supplementary Fig. S6: Impact of Netrin-1 inhibition on blood vessels and macrophages.**

**a-b**, Representative images of IHC staining for CD31 in cerulein-induced metaplastic pancreatic lesions from mice treated with Iso-antibody (a) or αNetrin-1 (b). DAPI was used for nuclear staining. Scale bar: 50 µm.

**c**, Quantification of CD31 fluorescence labeling in cerulein-induced metaplastic pancreatic lesions (M) from mice treated with Iso-antibody (Iso) or αNetrin-1 (αNet). Data are presented as mean ± SEM. $n = 31$ (Iso) and 28 (αNet) ROIs from 3 mice per group. ns: $p > 0.05$ (Mann–Whitney test).

**d-e**, Representative images of IHC staining for F4/80 and CD163 in cerulein-induced metaplastic pancreatic lesions labeled with E-cadherin (Ecadh) from mice treated with Iso-antibody (d) or αNetrin-1 (e). DAPI was used for nuclear staining. Scale bar: 50 µm.

**f-h**, Quantification of F4/80 (f) and CD163 (g) fluorescence labeling in cerulein-induced metaplastic pancreatic lesions (M) from mice treated with Iso-antibody (Iso) or αNetrin-1 (αNet). Panel (h) shows the CD163/F4/80 ratio. Data are presented as mean ± SEM. $n = 18$ (Iso) and 18 (αNet) ROIs from 3 mice per group. ns: $p > 0.05$ (two-tailed t- test).

**i-j**, Representative images of IHC staining for CD206 and iNOS in cerulein-induced metaplastic pancreatic lesions from mice treated with Iso-antibody (d) or αNetrin-1 (e). DAPI was used for nuclear staining. Scale bar: 50 µm.

**k**, Quantification of CD206 fluorescence labeling in cerulein-induced metaplastic pancreatic lesions (M) from mice treated with Iso-antibody (Iso) or αNetrin-1 (αNet). Data are presented as mean ± SEM. $n = 18$ (Iso) and 20 (αNet) ROIs from 3 mice per group. ns: $p > 0.05$ (2 tailed t- test).

**l**, Quantification of iNOS fluorescence labeling in cerulein-induced metaplastic pancreatic lesions (M) from mice treated with Iso-antibody (Iso) or αNetrin-1 (αNet). Data are presented as mean ± SEM. $n = 18$ (Iso) and 18 (αNet) ROIs from 3 mice per group. ns: $p > 0.05$ (Mann–Whitney test).

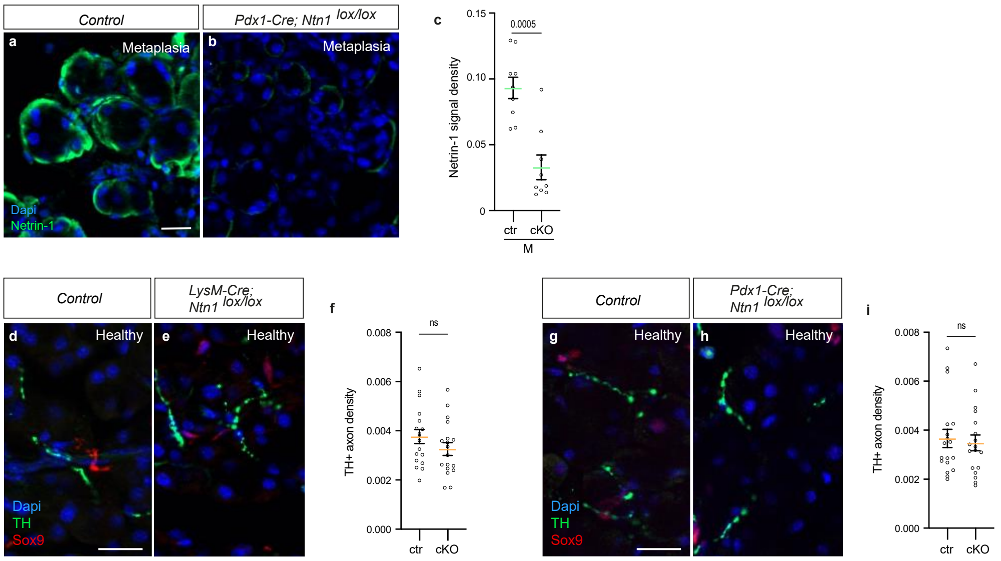


**Supplementary Fig. S7: Conditional deletion of *Ntn1* does not affect normal sympathetic innervation of the exocrine pancreas.**

**a-b,** Representative images of immunohistochemical (IHC) staining for Netrin-1 in the exocrine pancreas of the control (a) and *Pdx1-Cre–Ntn1*$^{lox/lox}$ mice (b) treated with cerulein. DAPI was used for nuclear staining. Scale bar: 20 µm.

**c**, Quantification of Netrin-1 fluorescence labeling in metaplastic lesions (M) of the exocrine pancreas of control (ctr) and *Pdx1-Cre–Ntn1*$^{lox/lox}$ (cKO) mice treated with cerulein. Data are presented as mean ± SEM. $n = 9$ sections from 3 mice/genotype. Significant p-values are indicated on the graph (Mann–Whitney test).

**d-e,** Representative images of IHC staining for TH and *SOX9* in the exocrine pancreas of control (d) and *LysM-Cre–Ntn1*$^{lox/lox}$ mice (e). DAPI was used for nuclear staining. Scale bar: 25 µm.

**f,** Quantification of TH$^{+}$ sympathetic axon density in the exocrine pancreas of control (ctr) and *LysM-Cre–Ntn1*$^{lox/lox}$ (cKO) mice. Data are presented as mean ± SEM. $n = 18$ sections from 3 mice per genotype. ns: $p > 0.05$ (unpaired t-test).

**g-h,** Representative images of IHC staining for TH and Sox9 in the exocrine tissues of control (g) and *Pdx1-Cre–Ntn1*$^{lox/lox}$ mice (h). DAPI was used for nuclear staining. Scale bar: 25 µm.

**i,** Quantification of TH$^{+}$ sympathetic axon density in the exocrine pancreas of control (ctr) and *Pdx1-Cre–Ntn1*$^{lox/lox}$ (cKO) mice. Data are presented as mean ± SEM. $n = 18$ sections from 3 mice per genotype. ns: $p > 0.05$ (unpaired t-test).

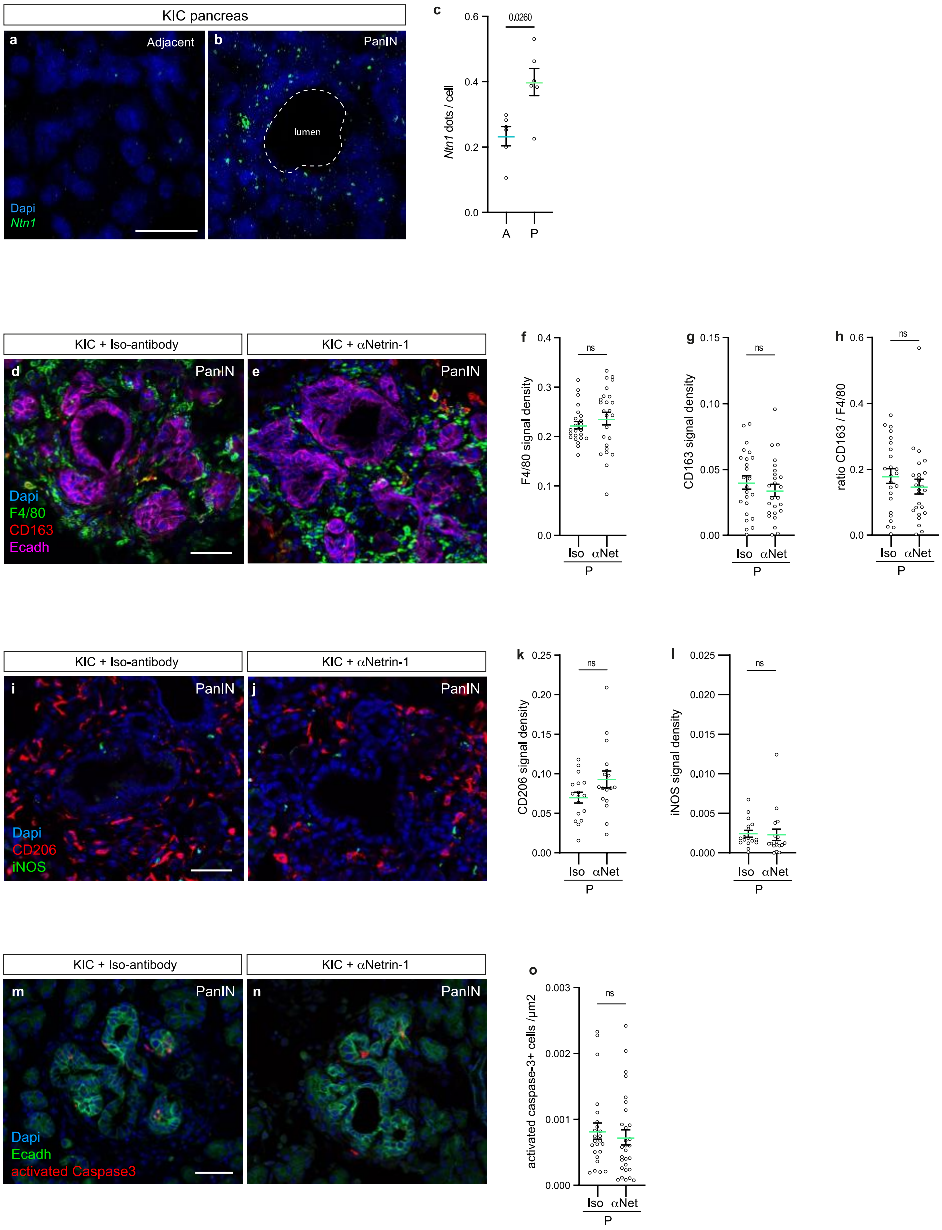


**Supplementary Fig. S8: Expression of Netrin-1 and effects on PanIN-associated macrophages and animal survival.**

**a-b**, Representative images of RNAscope *in situ* hybridization of *Ntn1* mRNA in pancreatic sections through PanIN lesions (b) and adjacent asymptomatic tissue (a) from KIC mice. DAPI was used for nuclear staining. Scale bar: 20 µm.

**c**, Quantification of *Ntn1* mRNA (dots) in PanIN (P) and adjacent (A) tissue from KIC mice. Data are presented as mean ± SEM. $n = 6$ PanINs and 6 asymptomatic ROIs from 3 KIC mice. p-value is indicated on the graph (Mann–Whitney test).

**d-e**, Representative images of IHC staining for F4/80 and CD163 in PanIN lesions labeled with E-cadherin (Ecadh) from KIC mice treated with Iso-antibody (d) or αNetrin-1 (e). DAPI was used for nuclear staining. Scale bar: 50 µm.

**f-h**, Quantification of F4/80 (f) and CD163 (g) fluorescence labeling in PanIN (P) from KIC mice treated with Iso-antibody (Iso) or αNetrin-1 (αNet). Panel (h) shows the CD163/F4/80 ratio. Data are presented as mean ± SEM. $n = 24$ (Iso) and 25 (αNet) ROIs from 3 KIC mice per group. ns: $p > 0.05$ (c and e: 2 tailed t- test; d: Mann–Whitney test).

**i-j**, Representative images of IHC staining for CD206 and iNOS in PanIN lesions labeled from KIC mice treated with Iso-antibody (i) or αNetrin-1 (j). DAPI was used for nuclear staining. Scale bar: 50 µm.

**k**, Quantification of CD206 fluorescence labeling in PanIN (P) from KIC mice treated with Iso-antibody (Iso) or αNetrin-1 (αNet). Data are presented as mean ± SEM. $n = 17$ (Iso) and 17 (αNet) ROIs from 3 KIC mice per group. ns: $p > 0.05$ (2 tailed t- test).

**l**, Quantification of iNOS fluorescence labeling in PanIN (P) from KIC mice treated with Iso-antibody (Iso) or αNetrin-1 (αNet). Data are presented as mean ± SEM. ($n = 17$ (Iso) and 17 (αNet) ROIs from 3 KIC mice per group. ns: $p > 0.05$ (Mann–Whitney test).

**m-n**, Representative images of IHC staining for activated caspase-3 in PanIN lesions labeled with E-cadherin (Ecadh) from KIC mice treated with Iso-antibody (m) or αNetrin-1 (n). DAPI was used for nuclear staining. Scale bar: 50 µm.

**o**, Quantification of activated caspase-3 fluorescence labeling in PanIN (P) from KIC mice treated with Iso-antibody (Iso) or αNetrin-1 (αNet). Data are presented as mean ± SEM. $n = 24$ (Iso) and 29 (αNet) ROIs from 3 KIC mice per group. ns: $p > 0.05$ (Mann–Whitney test).

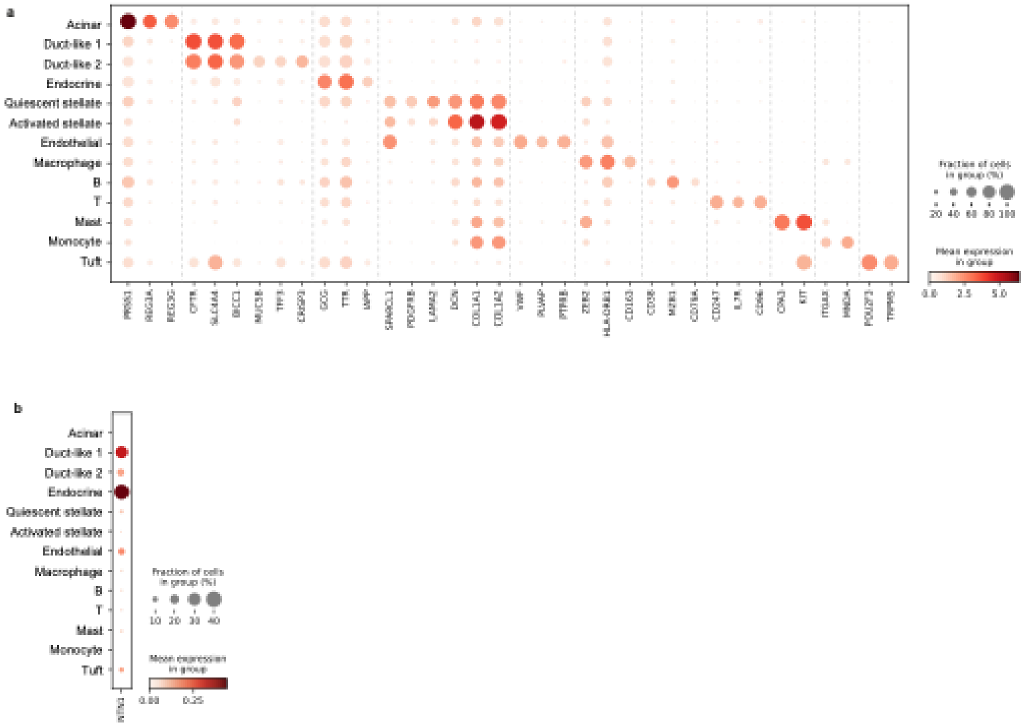


**Supplementary Fig. S9: Analysis of human chronic pancreatitis samples and NTN1 expression.**

**a**, Dot plot showing the expression of specific markers for the major cell types identified from single-nucleus RNA sequencing of human chronic pancreatitis samples.
**b**, Dot plot of NTN1 expression across all cell clusters.
The size of the dot represents the percentage of cells expressing a given gene, while the color indicates the average expression level.

### Supplementary Tables

#### Supplementary Table 1: Antibodies used in in vivo experiments

| Antibody name | Host | Dose | Supplier | Identifier |
|---|---|---|---|---|
| **Netrin-1 blocking antibody ($\alpha$Netrin-1)** | Mouse | 10 mg/kg | Netris Pharma Lyon, France | Cat# NP137, RRID AB_2811180 |
| **Isotypic control antibody (Iso-antibody)** | Mouse | 10 mg/kg | Netris Pharma Lyon, France | Cat# NP001 |

#### Supplementary Table 2: Primary antibodies

| Antibody name | Host | Dilution | Supplier | Identifier |
|---|---|---|---|---|
| **Anti-CGRP antibody** | Rabbit | 1:400 | Millipore | Cat# PC205L RRID AB_564312 |
| **Anti-CGRP antibody** | Goat | 1:200 | Abcam | Cat# ab36001 RRID AB_ 725807 |
| **Anti-TH antibody** | Chicken | 1:200 | Aves Labs | Cat # TYH RRID AB_10013440 |
| **Anti-TH antibody** | Rabbit | 1:200 | Millipore | Cat # AB152 RRID AB_390204 |
| **Anti-CK-19 antibody** | Rat | 1:40 | DSHB | Cat# TROMA-III RRID AB_2133570 |
| **Anti-CD31 antibody** | Rat | 1:400 | BD BioScience | Cat # 553370 RRID AB_394816 |
| **Anti-F4/80 antibody** | Rat | 1:200 | Thermo Fisher Scientific | Cat # 14-4801-85 RRID AB_467559 |
| **Anti-Iba1 antibody** | Rabbit | 1:500 | Antibodies-online | Cat # ABIN2857032 RRID AB_3096976 |
| **Anti-Sox9 antibody** | Rabbit | 1:1000 | Millipore | Cat # AB5535 RRID AB_2239761 |
| **Anti-Netrin-1 antibody** Mouse tissues | Goat | 1:500 | Abcam | Cat # ab122903 RRID AB_10898797 |
| **Anti-Netrin-1 antibody** Human tissues | Rabbit | 1:500 | Abcam | Cat # ab126729 RRID AB_11131145 |
| **Anti-DCC antibody** | Goat | 1:10 | R&D systems | Cat# AF844 RRID AB_2089765 |
| **Anti-UNC5B antibody** | Mouse | 1:500 | Abcam | Cat# ab54430 RRID AB_883377 |
| **Anti-VACht antibody** | Guinea pig | 1:200 | Synatic systems | Cat# 139105 RRID AB_10893979 |
| **Anti-VACht antibody** | Rabbit | 1:500 | Synatic systems | Cat# 139103 RRID AB_887864 |

| | | | | |
|---|---|---|---|---|
| **Anti-activated caspase3 antibody** | rabbit | 1:500 | Cell signalling | Cat# 9661L RRID AB_2341188 |
| **Anti- CD206 antibody** | Rat | 1:300 | BioLegend | Cat# 141707 RRID AB_10896057 |
| **Anti-CD163 antibody** | Rabbit | 1:100 | Non commercial | [1] |
| **Anti-Ecadherin antibody** | mouse | 1:500 | BD Biosciences | Cat # 610181 RRID AB_397581 |
| **Anti-KI67 antibody** | rat | 1:200 | BioLegend | Cat# 652402 RRID AB_11204254 |
| **Anti-iNOS antibody** | mouse | 1:100 | BD Biosciences | Cat# 610329 RRID AB_397718 |

**Supplementary Table 3: Secondary antibodies**

| Antibody name | Host | Dilution | Supplier | Identifier |
|---|---|---|---|---|
| **AF 488-anti-Goat IgG (H+L) antibody** | Donkey | 1:500 | Thermo Fisher Scientific | Cat# A11055 RRID AB_2534102 |
| **AF 568-anti-Goat IgG (H+L) antibody** | Donkey | 1:500 | Thermo Fisher Scientific | Cat# A11057 RRID AB_2534104 |
| **AF 647-anti-Goat IgG (H+L) antibody** | Donkey | 1:500 | Thermo Fisher Scientific | Cat# A-21447, RRID AB_2535864 |
| **AF 488-anti-Rabbit IgG (H+L) antibody** | Donkey | 1:500 | Thermo Fisher Scientific | Cat# A-21206 RRID AB_2535792 |
| **AF 568-anti-Rabbit IgG (H+L) antibody** | Donkey | 1:500 | Thermo Fisher Scientific | Cat#A10042, RRID: AB_2534017 |
| **AF 647-anti-Rabbit IgG (H+L) antibody** | Donkey | 1:500 | Thermo Fisher Scientific | Cat# A31573 RRID AB_2536183 |
| **AF 488-anti-Rat IgG (H+L) antibody** | Donkey | 1:500 | Thermo Fisher Scientific | Cat# A-21208 AB_141709 |
| **AF 568-anti-Rat IgG (H+L) antibody** | Donkey | 1:500 | Jackson Immuno Research Labs | Cat# 712-165-153 RRID AB_2340667 |
| **AF 647-anti-Rat IgG (H+L) antibody** | Donkey | 1:500 | Jackson Immuno Research Labs | Cat#712-605-153 RRID: AB_2340694 |
| **AF 488-anti-Mouse IgG (H+L) antibody** | Donkey | 1:500 | Thermo Fisher Scientific | Cat# A-21202 RRID AB_141607 |
| **AF 568-anti-Mouse IgG (H+L) antibody** | Donkey | 1:500 | Thermo Fisher Scientific | Cat# A10037 RRID AB_2534013 |
| **AF 647-anti-Mouse IgG (H+L) antibody** | Donkey | 1:500 | Jackson Immuno Research Labs | Cat# 715-605-151 RRID AB_2340863 |
| **AF 488-anti-Mouse IgG2a antibody** | Goat | 1:500 | Thermo Fisher Scientific | Cat# A-21131 RRID AB_2535771 |
| **AF 568-anti-Mouse IgG2a antibody** | Goat | 1:500 | Thermo Fisher Scientific | Cat# A-21134 RRID AB_2535773 |

| | | | | |
|---|---|---|---|---|
| **AF 647-anti-Mouse IgG2a antibody** | Goat | 1:500 | Thermo Fisher Scientific | Cat# A-21241 RRID AB_2535810 |
| **AF 488-anti-Chicken IgG (H+L) antibody** | Donkey | 1:500 | Jackson Immuno Research Labs | Cat# 703-545-155 RRID AB_2340375 |
| **AF 568-anti-Chicken IgG (H+L) antibody** | Donkey | 1:500 | Jackson Immuno Research Labs | Cat# 703 165 155 RRID AB_2340363 |
| **AF 647-anti-Chicken IgG (H+L) antibody** | Donkey | 1:500 | Jackson Immuno Research Labs | Cat# 703-605-155 RRID AB_2340379 |
| **AF 790-anti-Rat IgG (H+L) antibody** | Donkey | 1:500 | Jackson Immuno Research Labs | Cat# 712-655-153 RRID AB_2340701 |
| **anti-Rabbit HQ** | N/A | N/A | Roche | Cat# 760-4815 RRID AB_2811171 |
| **anti-HQ HRP** | N/A | N/A | Roche | Cat# 760-4820 RRID AB_3068525 |

### Supplementary Table 4: RNAscope probes

| Probe | Nb of pairs | Target region | Cat # |
|---|---|---|---|
| **RNAscope™ Probe- Mm-Ntn1-C2** | 20 | 3537 – 4505 NM_008744.2 | 407621-C2 |
| **RNAscope™ Probe- Mm-Iba1-C3** | 18 | 31 – 866 NM_019467.2 | 319141-C3 |
| **RNAscope™ Probe- Mm-Dcc-C3** | 20 | 644 – 1583 NM_007831.3 | 427491-C3 |
| **RNAscope™ Probe- Mm-Th** | 20 | 483 – 1603 NM_009377.1 | 317621 |
